# The hypoxic response extends lifespan through a bioaminergic and peptidergic neural circuit

**DOI:** 10.1101/2025.05.04.652087

**Authors:** Elizabeth S. Kitto, Shijiao Huang, Mira Bhandari, Cassie Tian, Rebecca L. Cox, Safa Beydoun, Emily Wang, Danielle Shave, Hillary A. Miller, Sarah A. Easow, Ella Henry, Eugene Chung, Megan L. Schaller, Scott F. Leiser

**Author notes:** Indicates co-first authors (Elizabeth S. Kitto, Shijiao Huang).

## Abstract

A coordinated response to stress is crucial for promoting the short- and long-term health of an organism. The perception of stress, frequently through the nervous system, can lead to physiological changes that are fundamental to maintaining homeostasis. Activating the response to low oxygen, or hypoxia, extends healthspan and lifespan in *C. elegans*. However, despite some positive impacts, negative effects of the hypoxic response in specific tissues prevent translation of their benefits in mammals. Thus, it is imperative to identify which components of this response promote longevity. Here, we interrogate the cell-nonautonomous signaling pathway downstream of genetic activation of the hypoxic response. We find that HIF-1-mediated signaling in ADF serotonergic neurons is both necessary and sufficient for lifespan extension. Signaling through the serotonin receptor SER-7 in the GABAergic RIS interneurons is necessary in this process. Our findings also highlight the involvement of additional neural signaling molecules, including the neurotransmitters tyramine and GABA, and the neuropeptide NLP-17, in mediating longevity effects. Finally, we demonstrate that oxygen- and carbon-dioxide-sensing neurons act downstream of HIF-1 in this circuit. Together, these insights develop a circuit for how genetic induction of the hypoxic response cell-nonautonomously modulates aging and suggests valuable targets for modulating aging in mammals.

## Introduction

The global population is aging at an unprecedented rate, posing significant challenges for healthcare systems worldwide^1^. The increasing burden of age-related diseases such as cardiovascular disease, diabetes, and neurodegeneration^2^ highlights the pressing need to improve healthspan, defined as the disease-free period of life^3^. Work over the past several decades has led to a better understanding of the conserved molecular and cellular mechanisms that regulate lifespan. One promising avenue has been the identification of environmental stressors that, when applied in a controlled manner, activate stress-response pathways to promote longevity. These interventions, including calorie restriction, exercise, hypoxia, heat shock, and cold exposure, extend lifespan and healthspan^4–7^. These stress-response pathways induce hormesis, a phenomenon in which a low dose of a stressor results in adaptive beneficial effects on cellular function^8,9^. Hormesis and the underlying stress-responses that promote longevity converge on cellular pathways, including oxidative stress, insulin signaling, autophagy, and protein homeostasis^10^, to promote longevity.

Multicellular organisms secrete a highly conserved set of neuromodulators including neurotransmitter and neuropeptide signals from the central nervous system. These signals integrate information about the organism’s external environment and internal state, and coordinate a physiological response in peripheral tissues. Within the context of aging, bioamine neurotransmitters contribute to the activation of many well-studied stress response mechanisms that promote longevity. For example, serotonin signaling modulates the mitochondrial unfolded protein response in *C. elegans*^11^, and regulates how food perception contributes to dietary restriction-mediated longevity in worms^12^ and flies^13^. Dopamine also modifies the longevity benefits of dietary restriction in invertebrates^12,14^, and polymorphisms in the dopamine D4 receptor are associated with longevity in humans^15^. Finally, adrenaline and its invertebrate analog, tyramine, have been linked to changes in the aging rate in both nematodes^14,16–18^ and mammals^19^. Hormone signals such as insulin^20–22^, growth factors^23,24^, and GnRH^14,25^ also contribute to longevity pathways across taxa. This growing body of work has led to great interest in manipulating longevity-promoting neural circuits to improve health^26^. While significant research effort focuses on pathways like autophagy, insulin signaling, and proteostasis in the context of aging, the role of hypoxia remains relatively underexplored. Currently, no longevity-promoting agents have been reported to target the hypoxic response.

The hypoxic response is highly conserved across species^27–29^, highlighting the relevance of this pathway in other organisms, including humans. In *C. elegans*, genetically activating the hypoxic response by knocking down or mutating the E3 ubiquitin ligase von hippel-lindau-1 (VHL-1)^30^ protein or stabilizing the hypoxia-inducible factor-1 (HIF-1) transcription factor^6,31^ is sufficient to extend lifespan. All of these interventions—environmental hypoxia, VHL-1 knockdown, and HIF-1 stabilization—require the intestinal enzyme flavin-containing monooxygenase-2 (*fmo-2)* to extend lifespan in *C. elegans*^31^. Hypoxic conditions also extend lifespan in a short-lived progeria mouse model^32^, and epidemiological studies indicate a correlation between hypoxia exposure and longevity^33,34^. However, the physiological changes induced by the hypoxic response are broad and involve adaptations such as increased vascularization, metabolic rewiring, and changes in cell survival pathways. In mammals, some of these same adaptations can be detrimental, as mutations in components of the hypoxic response have been linked to conditions like cancer and cardiovascular disease^35–38^. Additionally, HIF activity is essential to promote some forms of stress resistance to infection^39,40^ and oxidative stressors^41,42^ but can have detrimental effects on proteostasis^43^. This presents a key challenge: while hypoxia-induced longevity in a post-mitotic invertebrate model like *C. elegans* is promising, some of the mechanisms involved may not be directly translatable to mammals without causing deleterious side effects. Therefore, a mechanistic understanding of the individual cells, neural circuits, and pathways that mediate the beneficial but not detrimental effects of hypoxia on aging is essential to determine whether this circuit could be leveraged to improve human health.

Our previous work in *C. elegans* identified that stabilization of HIF-1 in neurons is sufficient to extend lifespan through the serotonin receptor, SER-7. This pathway eventually leads to the induction of *fmo-2*, a longevity gene expressed in the intestine^31,44^. In this study, we uncover key neural components of the longevity circuit initiated by genetic induction of the hypoxic response. Within this circuit, we identify individual cells, signals, and receptors necessary and/or sufficient to extend lifespan downstream of genetic activation of the hypoxic response. More specifically, we find serotonin signaling in the ADF serotonergic neurons is both necessary and sufficient to extend lifespan through genetic activation of the hypoxic response. This pathway signals through the serotonin receptor SER-7 in the RIS interneuron. We further demonstrate additional neurotransmitters (GABA and tyramine), and a neuropeptide (NLP-17) are critical for mediating these longevity effects. Finally, we identify that oxygen sensing neurons (URX, AQR, PQR and BAG) act downstream of neuronal HIF-1 in this circuit. Our insights into this longevity pathway provide a mechanistic understanding of how genetic activation of the hypoxic response delays aging and improves health.

## Results

### Serotonin signaling through the ADF neuron and the SER-7 receptor are necessary and sufficient for genetic activation of the hypoxic response to extend lifespan

Induction of the hypoxic response by targeted genetic manipulations can increase lifespan in *C. elegans.* These manipulations include decreasing activity of the VHL-1 E3 ubiquitin ligase that targets the transcription factor HIF-1 for degradation under hypoxia^30^ or by a mutation that stabilizes HIF-1 (HIF-1^P621A^)^6,31^ (**Fig. 1A**). We previously found that stabilizing HIF-1 in serotonergic neurons is sufficient to extend lifespan^31^. To determine which serotonergic neurons initiate HIF-1-mediated longevity, we generated strains with a nondegradable HIF-1 variant (HIF-1^P621A^)^31,45^ expressed under promoters specific to each of *C. elegans’* three primary serotonergic neuron types—the ADF, NSM, and HSN neurons^46^. These transgenic worms were then crossed into the *hif-1* null background to ensure that HIF stabilization in one neuron type could not feedback to modify HIF-1 activity in other cells. When we measured the lifespan of each strain relative to WT and *hif-1* knockout controls, we observed that stabilizing HIF-1 in the ADF or NSM neurons significantly extended lifespan by 26% and 23%, respectively (**Fig. 1B-C**). HIF-1 stabilization in the HSN neurons had a smaller effect but significantly extended lifespan by 9% (**Fig. 1D**). This result indicates that modifying signaling in any serotonergic neuron is sufficient to induce some level of hypoxic response-mediated longevity. These data suggest that serotonergic neurons can partially substitute for each other to promote longevity in this pathway. In summary, the ADF and/or NSM serotonergic neurons play an essential role in longevity following genetic activation of the hypoxic response.

**Figure 1.**
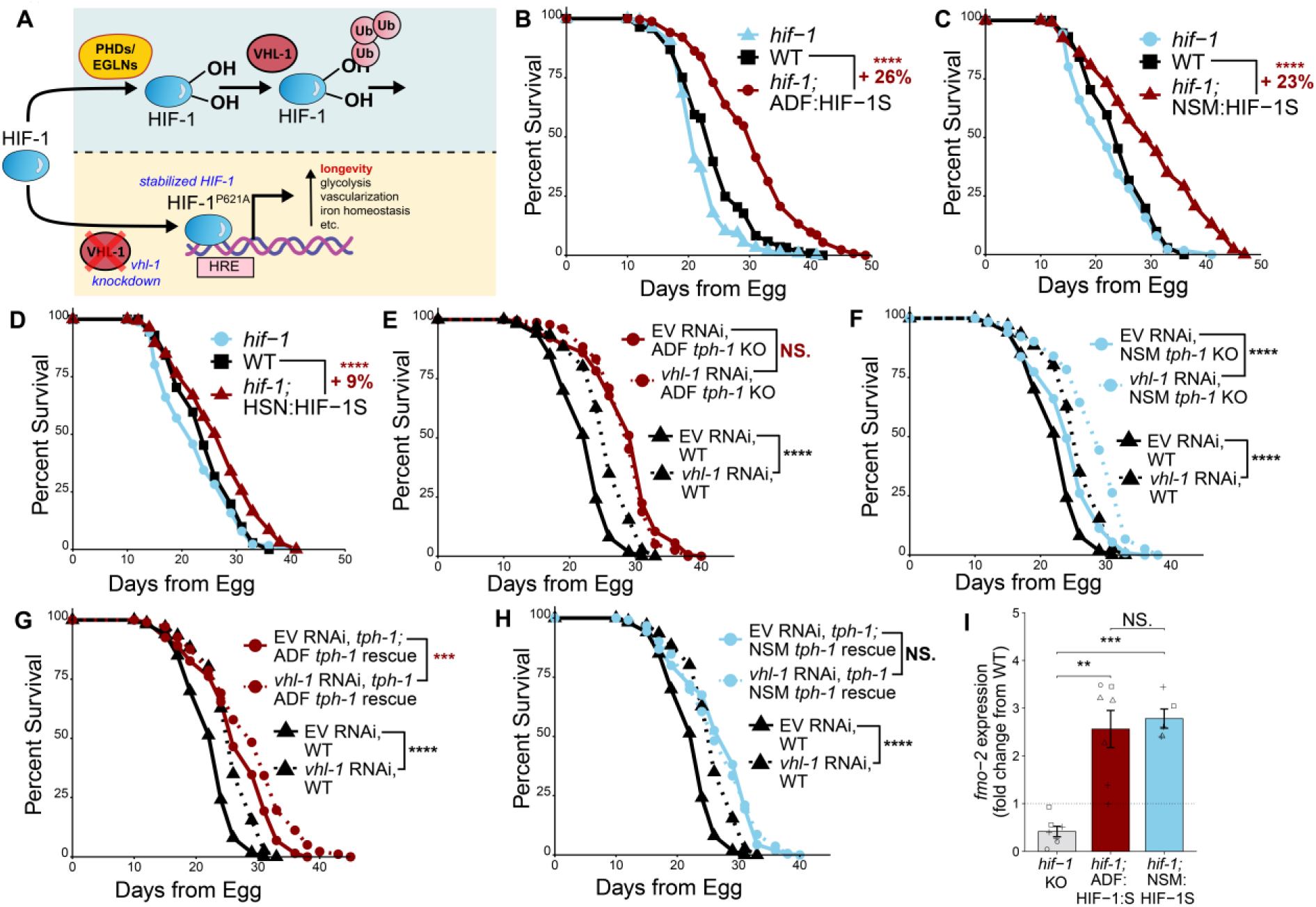
ADF serotonergic neurons are necessary and sufficient to extend lifespan downstream of genetic activation of the hypoxic response. (**A**) Diagram of the conserved hypoxic response and genetic approaches to activate it, *vhl-1* knockdown and HIF-1 stabilization. (**B-D**) Survival curves of WT, *hif-1(ia4)* knockout, *hif-1(ia4);* ADF::HIF-1S (**B**), *hif-1(ia4);* NSM::HIF-1S (**C**), and *hif-1(ia4);* HSN::HIF-1S (**D**) worms. *N* ≥ 256 (B), *N* ≥ 188 (**C**), and *N* ≥ 188 (**D**) worms per condition. (**E**) Survival curves of WT and ADF:*tph-1* knockout worms on empty vector (EV) or *vhl-1* RNAi. *N* ≥ 143 worms per condition. (**F**) Survival curves of WT and NSM:*tph-1* knockout worms on empty vector (EV) or *vhl-1* RNAi. *N* ≥ 193 worms per condition. (**G**) Survival curves of WT and *tph-1(mg280);* ADF:*tph-1* rescue worms on empty vector (EV) or *vhl-1* RNAi. *N* ≥ 178 worms per condition. (**H**) Survival curves of WT and *tph-1(mg280);* NSM:*tph-1* rescue worms on empty vector (EV) or *vhl-1* RNAi. *N* ≥ 160 worms per condition. (**I**) Gene expression of *fmo-2* in WT, *hif-1(ia4)* knockout, *hif-1(ia4);* ADF:HIF-1S, and *hif-1(ia4);* NSM:HIF-1S worms. Significance in panels B-H is from a log-rank test comparing median survival. Significance in panel I is from a Student’s t-test ^47^. In all panels, three replicates were plotted together, and all statistics include a Bonferroni correction for multiple comparisons. NS. = *p >* 0.05, * = *p* < 0.05, ** = *p* < 0.01, *** = *p* < 0.001, and **** = *p* < 0.0001.

After identifying this role for ADF and NSM HIF-1 stabilization, we also asked whether genetically activating the hypoxic response also affected healthspan in three different measures of *C. elegans* mobility: pumping rate, thrashing rate, and movement speed. We found that both ADF- and NSM-specific HIF-1 stabilization had no effect on pumping rate in young (day 1 of adulthood) worms. In aged animals (day 12 of adulthood), however, NSM-, but not ADF-, specific HIF-1 stabilization rescued the pumping rate decline in hif-1 knockout compared to WT worms (**Fig. S1C**). Similarly, NSM-, but not ADF-, specific HIF-1 stabilization rescued the thrashing rate decline in the hif-1 knockout young and aged worms **(Fig. S1D**). ADF and NSM HIF-1 stabilization also had no effect on average or maximum movement speed at days 1 and 5 of adulthood (**Fig. S1E-F**). Together, these results indicate that genetic activation of the hypoxic response in NSM neurons but not in the ADF neurons could improve healthspan.

HIF-1 activity also interacts with multiple stress responses, including oxidative stress, proteotoxic stress, and infection^41,42 39,40,43^. To determine whether various components of this pathway affect both longevity and the response to other stressors, and given that serotonin signaling modulates the mitochondrial unfolded protein response (mt-UPR)^48^, we asked whether stabilizing HIF-1 in these serotonergic neurons may also affect the mt-UPR. We found that HIF-1 stabilization in either the ADF or the NSM decreased the expression of *hsp-6* (**Fig. S1G**), a downstream effector of the mt-UPR^49^. This could suggest either that the mt-UPR response is impaired in these worms, or that HIF-1 stabilization decreases proteotoxic stress leading to a lower basal level of *hsp-6*.

To further test the necessity of the most impactful (for lifespan) serotonergic neurons in genetic activation of the hypoxic response-mediated longevity, we used transgenic animals in which *tph-1*, the rate-limiting enzyme in serotonin synthesis, was knocked out exclusively in the ADF or the NSM serotonergic neurons^12,50^. We then measured the lifespans of these strains on empty vector (EV) and *vhl-1* RNAi to determine the necessity of ADF and NSM serotonergic signaling in *vhl-1*-mediated longevity. Successful RNAi knockdown was confirmed with qPCR validation (**Fig. S1A**). We found that serotonin synthesis in ADF but not NSM neurons was required for *vhl-1* knockdown to extend lifespan (**Fig. 1E-F**). This indicates that while HIF stabilization in multiple serotonergic neurons can extend lifespan, ADF serotonin signaling is required for lifespan extension in response to *vhl-1* knockdown.

To test if serotonin production in a single neuron type is sufficient for genetic activation of the hypoxic response to extend lifespan, we rescued *tph-1* expression in a *tph-1* null background using promoters specific to the ADF or the NSM neurons^12^. The lifespans of these strains on EV or *vhl-1* RNAi were then measured. As expected and consistent with our previous publication^31^, the lifespan of the *tph-1* knockout strain was not extended by *vhl-1* knockdown (**Fig. S1B).** Consistent with the ADF neuron’s requirement for *vhl-1*-mediated longevity (**Fig. 1E-F**), serotonin synthesis in the ADF neurons was sufficient for *vhl-1* knockdown to extend lifespan while NSM serotonin synthesis was not (**Fig. 1G-H**). The enzyme *fmo-2* is induced in the intestine under hypoxia and is required for lifespan extension by environmental hypoxia, HIF-1 stabilization, and *vhl-1* depletion^31^. To test whether ADF- or NSM-specific HIF-1 stabilization also induces *fmo-2*, we measured *fmo-2* transcription in this strain using qPCR. We found that ADF and NSM HIF-1 stabilization significantly induced *fmo-2* expression relative to WT controls (**Fig. 1I**), suggesting a model in which ADF signaling extends lifespan through the same mechanism as *vhl-1* knockdown and environmental hypoxia. Together, these data indicate that while HIF-1 stabilization in any serotonergic neuron is sufficient to extend lifespan, only ADF serotonin production is necessary and sufficient for *vhl-1-*mediated longevity. This suggests that under normal physiological conditions the serotonergic ADF neurons propagate a signal in response to whole body stabilized HIF-1.

**Supplemental Figure 1.**
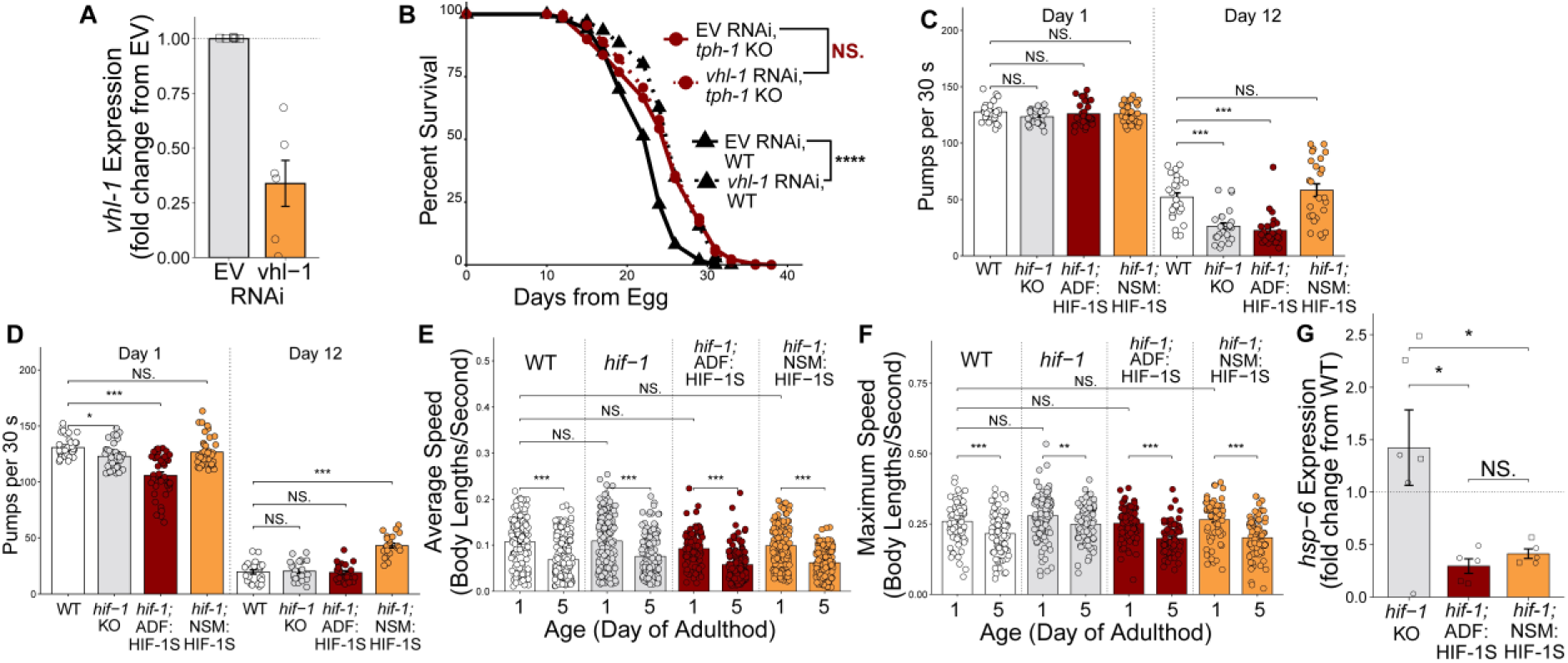
t*p*h*-1* is required for *vhl-1* mediated longevity. (**A**) Gene expression of *vhl-1* in worms on *vhl-1* RNAi compared to control worms on empty vector (EV) RNAi for two generations. *N* ≥ 200 worms per replicate, or 1200 worms per condition. The top of the bar represents the mean of the population and error bars indicate standard error of the mean (SEM). (**B**) Survival curve of WT, *vhl-1(ok161)*, *tph-1(mg280),* and *vhl-1(ok161); tph-1(mg280)* worms. *N* ≥ 338 worms per condition. Significance is from a log-rank test comparing median survival. (**C-D**) Quantification of the pumping (**C**) and thrashing (**D**) rates of WT, *hif-1(ia4)* knockout, *hif-1(ia4);* ADF::HIF-1S, and *hif-1(ia4);* NSM::HIF-1S worms at day 1, and 12 of adulthood. *N* ≥ 23 worms per condition. Significance is from a two-way ANOVA and post-hoc Tukey HSD test (unpaired, two-tailed). All panels show one representative replicate. Raw data from all three replicates can be found in supplemental data files. (**E-F**) The average (**E**) and maximum speed (**F**) of WT, *hif-1(ia4)* knockout, *hif-1(ia4);* ADF::HIF-1S, and *hif-1(ia4);* NSM::HIF-1S worms over a 30-minute video at days 1 and 5 of adulthood. *N ≥* 20 worms per condition. Significance is from a two-way ANOVA and post-hoc Tukey HSD test (unpaired, two-tailed). All panels show one representative replicate. Raw data from all three replicates can be found in supplemental data files. (**G**) Gene expression of *hsp-6* in *hif-1(ia4)* knockout, *hif-1(ia4);* ADF::HIF-1S, and *hif-1(ia4);* NSM::HIF-1S worms relative to WT controls. *N* ≥ 200 worms per replicate, or 1,000 worms per condition. Significance is from a two-sided Wilcoxon rank-sum test. In all bar plots, the top of the bar represents the mean of the population and error bars indicate standard error of the mean (SEM). In all panels, NS. = *p >* 0.05, * = *p* < 0.05, ** = *p* < 0.01, *** = *p* < 0.001, and **** = *p* < 0.0001. In panels A-B and G, three replicates were plotted together, and all statistics include a Bonferroni correction for multiple comparisons.

After identifying ADF neurons as the most central serotonergic neurons in longevity following genetic activation of the hypoxic response, we next sought to determine the downstream receptor responding to serotonin signaling from ADF neurons. *C. elegans* have six known serotonin receptors—SER-1, SER-4, SER-5, SER-7, MOD-1, and LGC-50. Of these six receptors, our previous findings showed that *ser-7* expression is required for *vhl-1* knockdown to induce *fmo-2* and extend lifespan^31^. SER-7 is a G protein-coupled receptor (GPCR) with high sequence identity to the 5-HT_7_ receptor in mammals^51,52^. *ser-7* is thought to be expressed in 27 of *C. elegans’* 302 neurons^53^ and contributes to many aspects of physiology such as reproduction and pharyngeal pumping^51^. Consequently, broad manipulation of SER-7 signaling is not an ideal method for exclusively modifying aging. To identify more specific targets for lifespan extension within this pathway, we asked whether *ser-7* expression in any subset of neurons is sufficient to rescue the *vhl-1* mediated *fmo-2* induction in a *ser-7* null background.

We created transgenic strains with *ser-7* expression under the control of 11 different cell-specific promoters in a *ser-7* null background. These promoters were selected to rescue *ser-7* expression in the following neuronal populations: whole-body rescue (*ser-7p::ser-7)*, interneuron rescue (*glr-1p::ser-7)*, bioaminergic neuron rescue (*cat-1p::ser-7)*, glutamatergic neuron rescue (*eat-4p::ser-7*), GABAergic neuron rescue (*unc-47p::ser-7*), cholinergic neuron rescue (*unc-17p::ser-7*), sensory neuron rescue (*osm-6p::ser-7*), GABAergic motor neuron rescue (*unc-25p::ser-7*), cholinergic motor neuron rescue (*acr-2p::ser-7*), M3 and M4 neuron rescue (*ceh-28p::ser-7*), and intestinal rescue (*vha-6p::ser-7*) in a *ser-7* null background. To measure whether *ser-7* expression restores *vhl-1* mediated longevity, we injected each construct into *ser-7* null worms crossed with a single-copy transcriptional reporter for *fmo-2 (fmo-2p::mCherry)*. Because *fmo-2* is induced by and required for hypoxia-mediated longevity^31^, we utilized its induction as an efficient screening tool to identify components of this longevity pathway.

We first confirmed that *ser-7* knockout attenuates *vhl-1-*mediated *fmo-2* induction (**Fig. 2A**). The whole-body rescue of *ser-7* expression under the endogenous *ser-7* promoter restored *fmo-2* induction to 91% of WT controls (**Fig. 2A**). Of the 10 cell-specific *ser-7* rescues, expression in interneurons, bioaminergic neurons, glutamatergic neurons, or GABAergic neurons fully rescued *vhl-1*-mediated *fmo-2* induction to ≥ 100% of the WT control response (**Fig. 2A**, pink bars). *ser-7* rescue in GABAergic motor neurons or in sensory neurons showed a partial rescue (defined as a 50-99% increase from *fmo-2* induction in the *ser-7* knockout) (**Fig. 2A**, orange bars). In contrast, rescuing *ser-7* in cholinergic motor neurons, cholinergic neurons, M3 and M4 neurons, or in the intestine did not rescue (**Fig. 2A**, gray bars). Taken together, 6 of the 10 cell-specific *ser-7* rescues at least partially restored *vhl-1-*mediated *fmo-2* induction, including some that have no known overlap in *ser-7* expression patterns. Collectively, these data suggest that serotonin receptor *ser-7* expression in multiple neurons may be sufficient to convey the hypoxic signal to the intestine and induce *fmo-2*.

**Figure 2.**
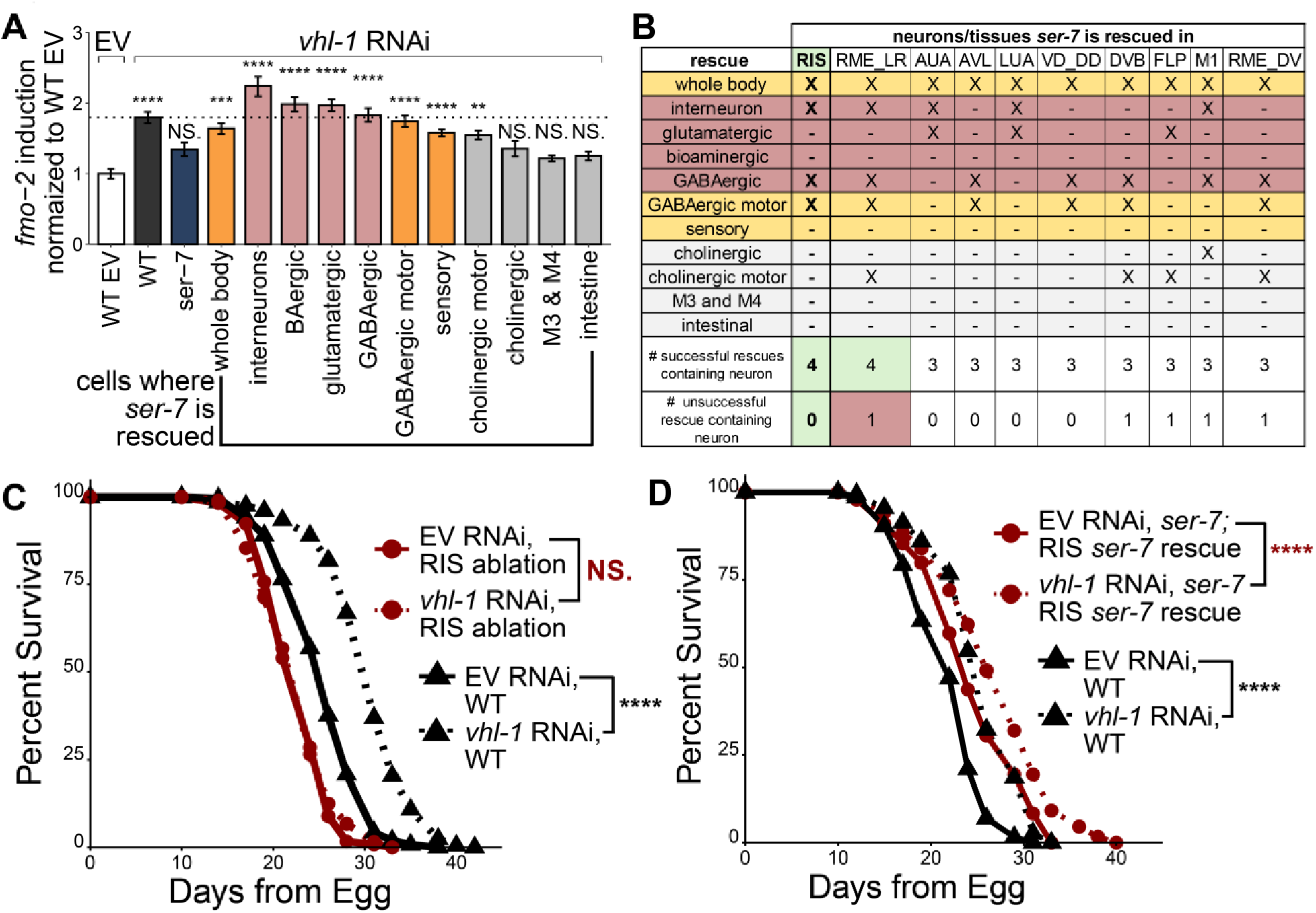
*ser-7* expression in the GABAergic RIS neuron is required for genetic activation of the hypoxic response to extend lifespan. (**A**) Quantification of *fmo-2p::mCherry* (WT positive control), *fmo-2p::mCherry; ser-7(tm1325)* knockout (negative control), and *fmo-2p::mCherry; ser-7(tm1325)* knockouts with *ser-7* rescued in the whole body (*ser-7p::ser-7)*, interneurons (*glr-1p::ser-7)*, bioaminergic neurons (*cat-1p::ser-7)*, glutamatergic neurons (*eat-4p::ser-7*), GABAergic neurons (*unc-47p::ser-7*), GABAergic motor neurons (*unc-25p::ser-7*), sensory neurons (*osm-6p::ser-7*), cholinergic motor neurons (*acr-2p::ser-7*), cholinergic neurons (*unc-17p::ser-7*), M3 & M4 neurons (*ceh-28p::ser-7*), and the intestine (*vha-6p::ser-7*) on empty vector (EV, WT control shown) or *vhl-1* RNAi (all strains shown). Bar height indicates the mean fluorescence on *vhl-1* RNAi of each genotype normalized to the empty vector control value from that genotype. The dashed line indicates the *vhl-1-*mediated *fmo-2* induction of WT positive control. *N* ≥ 45 worms per condition. Error bars indicate SEM. Significance is from a two-way ANOVA (*fmo-2* induction ∼ genotype*RNAi) and post-hoc Tukey HSD test ^47^. Stars signify results from this post-hoc test comparing the *fmo-2* induction of each strain on EV RNAi the *fmo-2* induction of that strain on *vhl-1* RNAi. (**B**) A table showing the 10 *ser-7* expressing candidate neurons and their expression patterns among the *ser-7* cell-specific strains that successfully rescued *fmo-2* induction in Fig. 2A. Pink rows indicate rescues that full restored *vhl-1* mediated *fmo-2* induction, orange rows indicate rescues that partially restored *vhl-1-*mediated *fmo-2* induction, and gray rows signify unsuccessful rescue constructs from the data in Fig. 2A. (**C**) Survival curve of WT and RIS ablation (*Ex[srsx-18p::caspase-3(p12)-nz]; Ex[srsx-18p::cz-caspase-3 (p17)]; srsx-18p::GFP*) worms on empty vector (EV) and *vhl-1* RNAi. *N* ≥ 192 worms per condition. (**D**) Survival curve of WT and *ser-7 (tm1325); ser-7* rescue in the RIS neuron (*flp-11p::ser-7*) worms on EV and *vhl-1* RNAi. *N* ≥ 144 worms per condition. Significance in panels C-D is from a log-rank test comparing median survival. In all panels, NS. = *p >* 0.05, * = *p* < 0.05, ** = *p* < 0.01, *** = *p* < 0.001, and **** = *p* < 0.0001. In all panels, three replicates were plotted together, and all statistics include a Bonferroni correction for multiple comparisons.

We compiled a table containing the top candidates for the *ser-7* neuron based on which rescue constructs restored *ser-7* expression (simplified table in **Fig. 2B**, full table in **Fig. S2A**). We hypothesized that neurons present in many of the successful rescue constructs would be most important in the *vhl-1-*mediated longevity circuit. Conversely, we hypothesized that candidate neurons within rescues that did not restore a WT-like response are less likely to be important (**Fig. 2B**; **Fig. S2A**). We acknowledge that this analysis strategy is biased toward neurons rescued in a greater proportion of the transgenic strains, which could lead to false negatives. From this analysis, we identified 10 neurons that were rescued in 3-4 of the 6 successful rescue constructs (**Fig. 2B**). Five of these neurons (DVB, FLP, M1, RME_LR, and RME_DV) were also present in unsuccessful rescue constructs, so they were not considered for follow-up experiments. Of the remaining five neurons (RIS, AUA, AVC, LUA, and VD_DD), the top candidate was the RIS neuron, because RIS was present in the highest number (4 as shown in **Fig. 2B**) of successful rescue constructs.

**Supplemental Figure 2.**
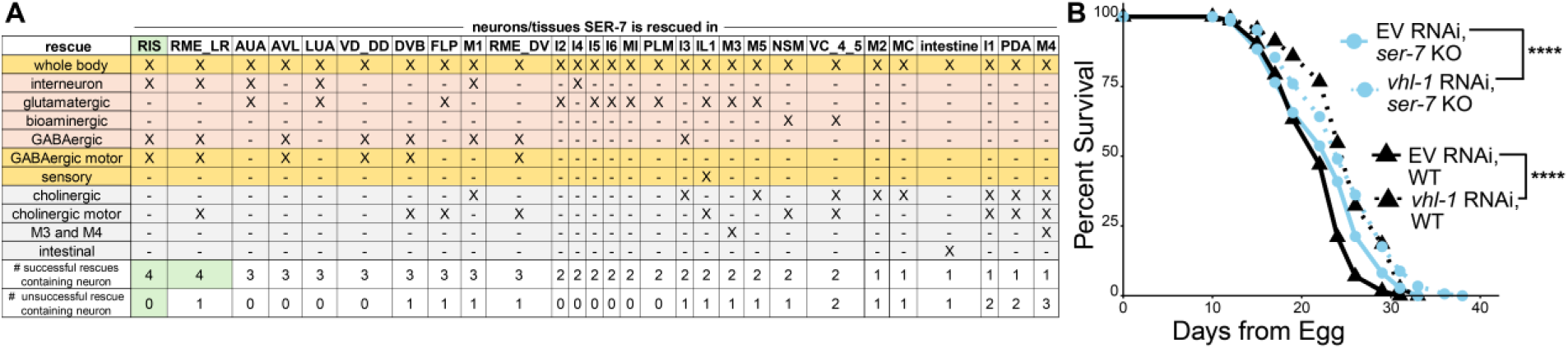
s*e*r*-7* expression in each *ser-7* expressing neuron is rescued by at least one construct, and *ser-7* is required for *vhl-1-*mediated longevity. (**A**) A table showing all *ser-7* expressing candidate neurons, and which constructs in Fig. 2A restored *ser-7* expression in these neurons. Pink rows indicate rescues that fully restored *vhl-1* mediated *fmo-2* induction, orange rows indicate rescues that partially restored *vhl-1-*mediated *fmo-2* induction, and gray rows signify unsuccessful rescue constructs from the data in panel A. (**B**) Survival curve of WT, and *ser-7(tm1325)* worms on empty vector (EV) and *vhl-1* RNAi. *N* ≥ 259 worms per condition. Significance is from a log-rank test comparing median survival. Cox Regression for an interaction between the effect of *ser-7 (tm1325)* knockout and *vhl-1* RNAi knockdown on lifespan. *p* = 0.003, **. NS. = *p >* 0.05, * = *p* < 0.05, ** = *p* < 0.01, *** = *p* < 0.001, and **** = *p* < 0.0001. Three replicates were plotted together, and all statistics include a Bonferroni correction for multiple comparisons.

To validate whether the RIS neuron is a key signaling cell in *vhl-1*-mediated longevity, we first genetically ablated the RIS neuron to test its necessity. Our results showed that *vhl-1* RNAi does not extend lifespan of RIS ablated worms, indicating that the RIS neuron is completely required for *vhl-1* knockdown to extend lifespan (**Fig. 2C**). To test the sufficiency of RIS signaling, we next rescued *ser-7* expression in the RIS neuron in the *ser-7* null background under the *flp-11* promoter. *flp-11* is expressed in the RIS neuron^54^ and uv1 neuroendocrine cells^55^ that do not endogenously express *ser-7*^53^. We first confirmed that *ser-7* knockout worms are partially required for *vhl-1* RNAi to extend lifespan (**Fig. S2B**, Cox Regression, *p* = 0.003, **). Significantly, *ser-7* rescue in the RIS neuron (*ser-7; flp-11p::ser-*7) restores the lifespan extension by *vhl-1* RNAi to the same degree as WT worms (**Fig. 2D**). These data suggest that *ser-7* expression in the RIS neuron only is sufficient for *vhl-1-*mediated longevity. However, one limitation of this approach is that non-physiological expression of *ser-7* in the uv1 cells could play a role in restoration of WT-like *vhl-1* mediated *fmo-2* induction. Together, these data are consistent with a model where genetic activation of the hypoxic response stabilizes HIF-1 in the ADF neurons, which modifies serotonin signaling to the SER-7 receptor on the RIS interneuron.

### Tyraminergic signaling from the RIM neuron and the tyramine receptor TYRA-3 are required for *vhl-1* knockdown to extend lifespan

After identifying neuron subtypes involved in serotonin signaling downstream of genetic activation of the hypoxic response, we next set out to identify whether other neurotransmitters act in the pathway. To answer this question, we obtained mutants deficient in the production of each type of neurotransmitter in *C. elegans* (**Fig. 3A**) and measured their lifespan on control and *vhl-1* RNAi. Out of the six neurotransmitters we tested, blocking the production of GABA (*unc-25*, **Fig. 3B**) or tyramine + octopamine (*tdc-1,* **Fig. 3C**) attenuated longevity on *vhl-1* RNAi. The requirement of GABA is consistent with our finding that *ser-7* expression on the GABAergic RIS neuron is also necessary for *vhl-1*-mediated longevity (**Fig. 2**), connecting the serotonergic and GABAergic components of this circuit.

**Figure 3.**
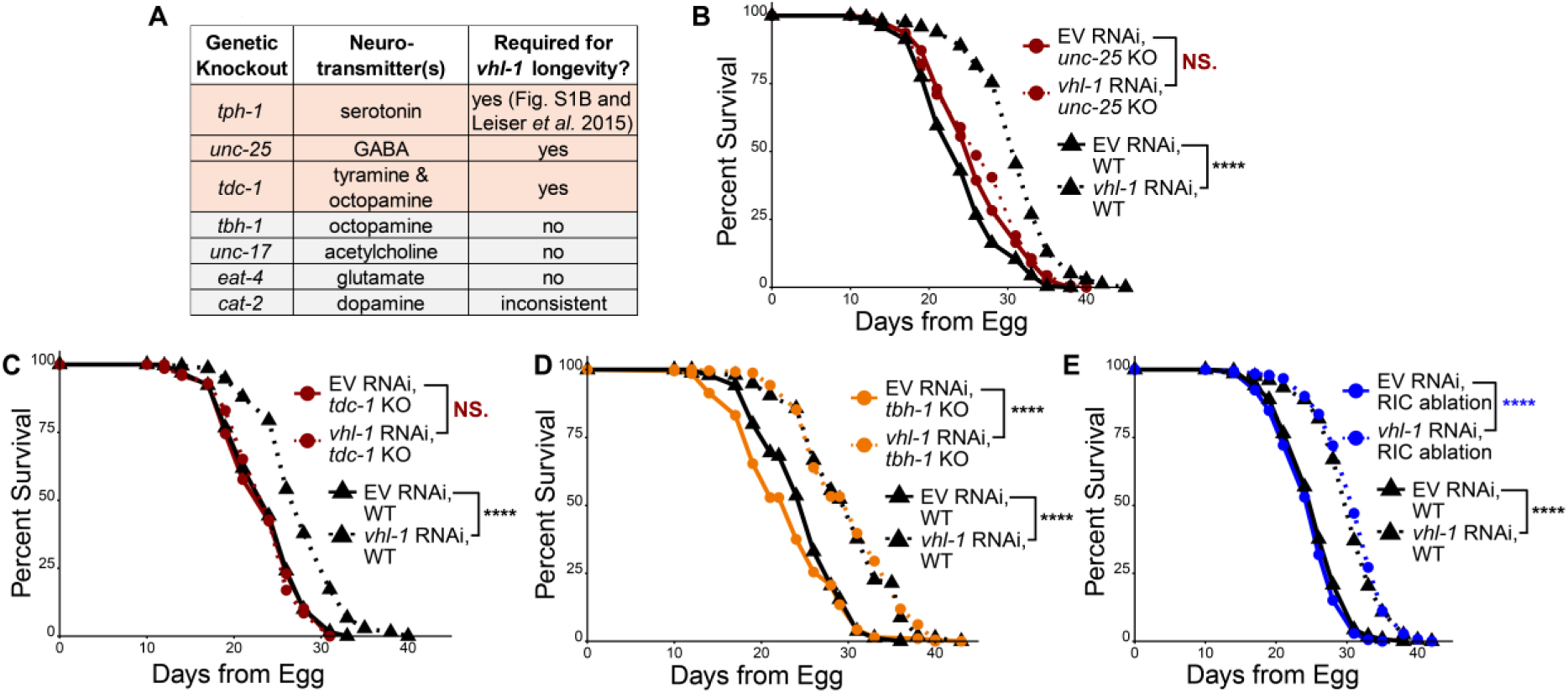
GABA and tyramine synthesis are required for *vhl-1* knockdown to extend lifespan. (**A**) Summary table of the necessity of each *C. elegans* neurotransmitter for *vhl-1* RNAi*-*mediated longevity. (**B-E**) Survival curves of WT and *unc-25(e156)* (GABA deficiency) (**B**), *tdc-1(n3419)* (tyramine + octopamine deficiency) (**C**), *tbh-1(n3247)* (octopamine deficiency) (**D**), and RIC ablation (*Ex[tbh-1p::caspase-3(p12)-nz]; Ex[tbh-1p::cz-caspase-3 (p17)]; tbh-1p::GFP*) (**E**) worms on empty vector (EV) and *vhl-1* RNAi. *N* ≥ 310 (**B**), ≥ 201 (**C**), ≥ 258 (**D**), and ≥ 258 (**E**) worms per condition. Significance in panels B-E is from a log-rank test comparing median survival. NS. = *p >* 0.05, * = *p* < 0.05, ** = *p* < 0.01, *** = *p* < 0.001, and **** = *p* < 0.0001. In all panels, three replicates were plotted together, and all statistics include a Bonferroni correction for multiple comparisons.

Tyramine (analogous to mammalian adrenaline^56^) is synthesized from tyrosine by tyrosine decarboxylase (*tdc-1*) and can be further converted to octopamine (analogous to mammalian noradrenaline) by tyramine β-hydroxylase (*tbh-1*) in *C. elegans*^57^. Therefore, the *tdc-1* knockout is deficient in both tyramine and octopamine synthesis (**Fig. 3C**) and the *tbh-1* knockout (**Fig. 3D**) is only deficient in octopamine synthesis. Since worms lacking only octopamine (*tbh-1*, **Fig. 3D**) were long-lived on *vhl-1* RNAi but worms lacking both tyramine and octopamine (*tdc-1*, **Fig. 3C**) were not, we can conclude that tyramine is required for genetic activation of the hypoxic response to extend lifespan. To further test whether tyramine but not octopamine is involved in this pathway, we generated a strain in which the primary octopamine-producing neuron, RIC^57^, is genetically ablated. We found that the RIC neuron is not required for *vhl-1-*mediated longevity (**Fig. 3E**), further supporting that tyramine but not octopamine is a key neurotransmitter in the circuit driving longevity after genetic activation of the hypoxic response.

In addition to octopamine (*tbh-1,* **Fig. S3C**), we also observed that acetylcholine (*unc-17*, **Fig. S3A**) and glutamate (*eat-4*, **Fig. S3B**) were not required for lifespan extension by *vhl-1* RNAi. Our lifespans of the dopamine synthesis mutant *cat-2* indicated that dopamine is inconsistently required for genetic activation of the hypoxic response to extend lifespan. In three biological replicates, we observed a full requirement, partial requirement, and no requirement for dopamine synthesis in *vhl-1* RNAi-mediated longevity. When analyzed together, the net result suggests that dopamine is partially required for *vhl-1-*mediated longevity (**Fig. S3C**, Cox Regression for interaction between genotype and *vhl-1* RNAi, *p <* 0.0001, ****). We did not follow up on dopamine due to the inconsistency. Together, the results from this neurotransmitter screen support the conclusion that in addition to serotonin, both GABA and tyramine play a role in *vhl-1*-mediated longevity.

**Supplemental Figure 3.**
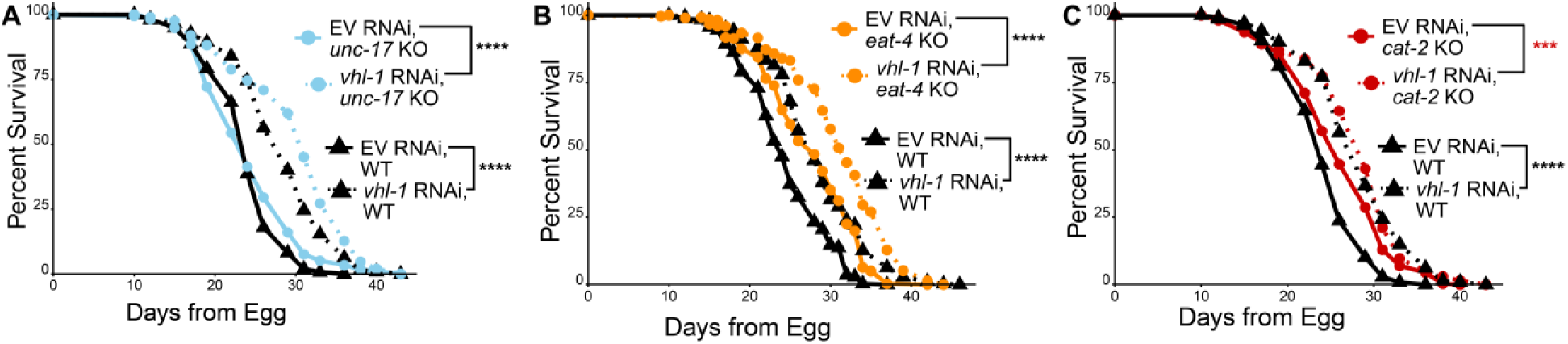
Acetylcholine, glutamate, and dopamine synthesis are not fully required for *vhl-1* RNAi to extend lifespan. (**A-C**) Survival curves of WT and *unc-17(e113)* (acetylcholine deficiency) (**A**), *eat-4(ky5)* (glutamate deficiency) (**B**), and *cat-2(n4547)* (dopamine deficiency) (**C**) worms on empty vector (EV) and *vhl-1* RNAi. *N* ≥ 213 (**A**), ≥ 274 (**B**), and ≥ 270 (**C**) worms per condition. (**C**) Cox Regression for an interaction between genotype and *vhl-1* RNAi, p < 0.0001, ****. Significance in all panels is from a log-rank test comparing median survival. NS. = *p >* 0.05, * = *p* < 0.05, ** = *p* < 0.01, *** = *p* < 0.001, and **** = *p* < 0.0001. In all panels, three replicates were plotted together, and all statistics include a Bonferroni correction for multiple comparisons.

Nematodes synthesize GABA in 26 of 302 neurons while tyramine is only synthesized in two cells^58^. As a result, identifying highly specific components of tyraminergic signaling is more feasible than mapping GABAergic signaling components. Therefore, we sought to determine the tyramine-producing neuron(s) and tyramine receptor(s) that contribute to *vhl-1*-mediated longevity. In *C. elegans*, the two canonically tyraminergic cell types (the RIM neuron and the uv1 neuroendocrine cells) express *tdc-1* but not *tbh-1*^58^. To determine which tyraminergic cell(s) act in the*vhl-1*-mediated longevity pathway, we expressed *tdc-1* under promoters specific to either RIM or uv1 in a *tdc-1* null background^14^. We then measured whether the lifespans of these tyramine rescue strains could be extended by *vhl-1* RNAi. Our results showed that *vhl-1* RNAi extended lifespan in the RIM rescue strain (**Fig. 4A**), but not in the uv1 rescue (**Fig. 4B**). This finding demonstrates that tyramine synthesis in the RIM neuron is sufficient for *vhl-1* RNAi to extend lifespan. Notably, both tyramine rescue constructs lived longer than WT controls (**Fig. 4A-B**). This could indicate that altering tyramine signaling modifies lifespan both within and independently of the *vhl-1-*mediated longevity pathway.

**Figure 4.**
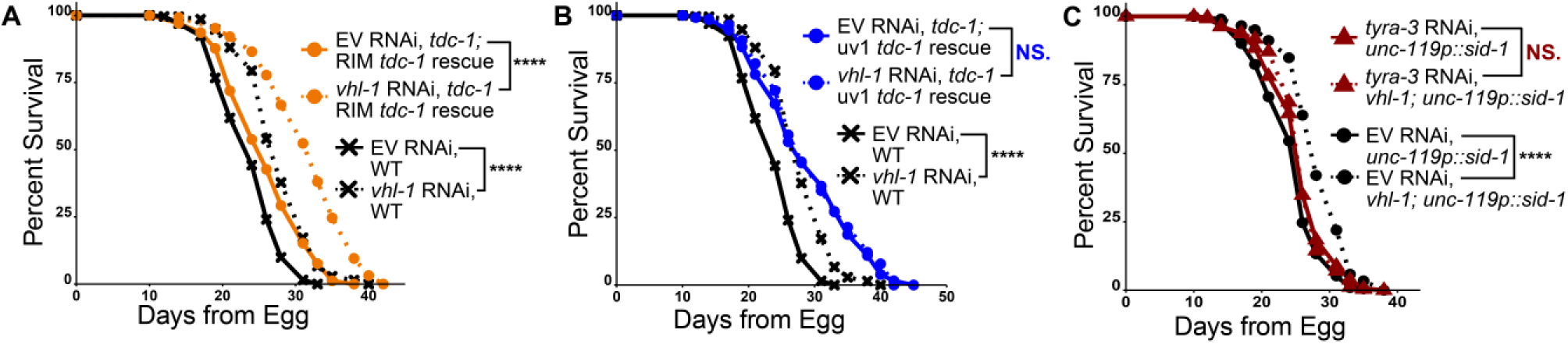
The RIM neuron and the tyramine receptor *tyra-3* act in the *vhl-1*-mediated longevity pathway. (**A-B**) Survival curves of WT, RIM rescue (*tdc-1 (n3419);* Ex[*ocr-4p::tdc-1])* (**A**), and uv1 *tdc-1* rescue (*tdc-1 (n3419);* Ex[*gcy-13p::tdc-1])* (**B**) strains on empty vector (EV) and *vhl-1* RNAi. *N* ≥ 126 (**A**) and *N* ≥ 140 (**B**) worms per condition. (**C**) Survival curves of TU3311(*unc-119p::sid-1)* and *vhl-1(ok161);* TU3311(*unc-119p::sid-1*) worms on empty vector (EV) and *tyra-3* RNAi. *N* ≥ 215 worms per condition. Significance in all panels is from a log-rank test comparing median survival. NS. = *p >* 0.05, * = *p* < 0.05, ** = *p* < 0.01, *** = *p* < 0.001, and **** = *p* < 0.0001. In all panels, three replicates were plotted together, and all statistics include a Bonferroni correction for multiple comparisons.

To identify the tyramine receptor(s) necessary for genetic activation of the hypoxic response to extend lifespan we used RNAi to knockdown each of *C. elegans’* four known tyramine receptors^59–62^ (**Fig. 4C**). Since tyramine receptors are mostly expressed in neurons, and RNAi uptake into some neurons is less efficient than other cell types, we used a strain with enhanced neuronal RNAi uptake (TU3311, *unc-119p::sid-1*) crossed into *vhl-1* knockout worms. This neural enhanced RNAi uptake strain was not used previously in this work for *vhl-1* RNAi experiments because *vhl-1* RNAi knockdown extends lifespan in WT worms^31^. The generated *vhl-1; unc-119p::sid-1* strain was long-lived compared to the control strain *unc-119p::sid-1* (**Fig. 4C**). Successful RNAi knockdown was validated via qPCR (**Fig. S4A**). Our lifespan results show that the tyramine receptor *tyra-3* was fully required for *vhl-1-*mediated longevity (**Fig. 4C**), but the tyramine receptors *tyra-2* and *ser-2* were not (**Fig. S4B-C**). Another tyramine receptor, *Igc-55,* showed a partial requirement for *vhl-1*-mediated longevity (**Fig. S4D**). Although there was a minor lifespan extension by *vhl-1* knockout on *Igc-55* RNAi (**Fig. S4D**), we did observe a significant interaction between *lgc-55* knockdown and genotype on lifespan (Cox Regression, *p* < 0.0001, ****), indicating a partial requirement. In summary, these data indicate that tyramine signaling from the RIM neuron to the TYRA-3 receptor is required for *vhl-1*-mediated longevity, while the LGC-55 receptor may play a partial role. Interestingly, TYRA-3 is highly expressed on the low oxygen/high carbon dioxide-sensing BAG neuron as well as in the intestine^53,63^, the site of *fmo-2* induction during hypoxia^31^.

**Supplemental Figure 4.**
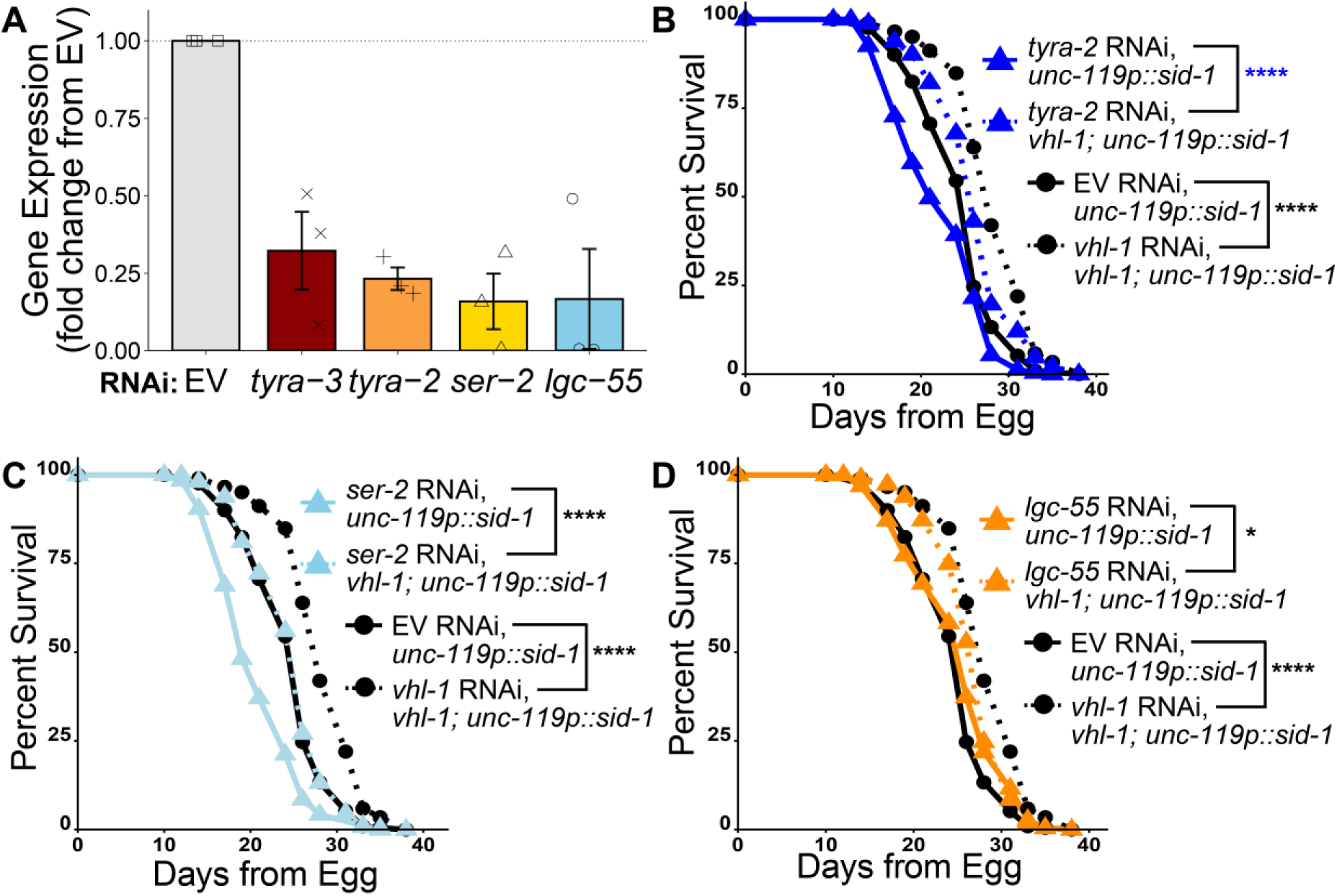
t*y*ra*-2* and *ser-2* are not necessary for *vhl-1-*mediated longevity, while *lgc-55* is partially required. (**A**) Gene expression of *tyra-3, tyra-2, ser-2,* and *lgc-55* in TU3311(*unc-119p::sid-1)* worms raised on RNAi targeting each gene compared to control worms raised on empty vector (EV) RNAi for two generations. *N* ≥ 300 worms per replicate, or 900 worms per condition. The top of the bar represents the mean of the population and error bars indicate standard error of the mean (SEM). (**B-D**) Survival curves of TU3311(*unc-119p::sid-1)* and *vhl-1(ok161);* TU3311(*unc-119p::sid-1*) worms on empty vector (EV) and *tyra-2* (**B**), *ser-2* (**C**), or *lgc-55* (**D**) RNAi. Cox Regression for an interaction between *lgc-55* knockdown and *vhl-1* knockout on lifespan in the TU3311background strain. *p* < 0.0001, ****. *N* ≥ 215 worms per condition. Significance is from a log-rank test comparing median survival. NS. = *p >* 0.05, * = *p* < 0.05, ** = *p* < 0.01, *** = *p* < 0.001, and **** = *p* < 0.0001. In all panels, three replicates were plotted together, and all statistics include a Bonferroni correction for multiple comparisons.

### Oxygen and carbon dioxide sensing neurons act downstream of serotonin signaling

Given that *tyra-3* is highly expressed in the canonical low-oxygen sensing neuron BAG^53^, we next explored whether neurons responsible for sensing high- and low-oxygen conditions are essential for genetic activation of the hypoxic response to extend lifespan. *C. elegans* have four oxygen sensory neurons: the URX, PQR, and AQR neurons detect high levels of oxygen^63^, the BAG neuron responds to low levels of oxygen^64^ and high levels of carbon dioxide^65^ (**Fig. 5A**). We hypothesized that these oxygen sensing cells may perceive hypoxic conditions and help initiate the hypoxic response. Alternatively, these oxygen sensing neurons could modify behavior in response to hypoxia but not play a role in longevity. We found that *vhl-1* knockdown was unable to extend lifespan when we genetically ablated the three high-oxygen sensing neurons (URX, AQR, PQR, **Fig. 5B**). Interestingly, the low O_2_/high CO_2_ responsive BAG neurons were also required for *vhl-1-*mediated longevity (**Fig. 5C**). To test the epistasis of oxygen sensing neurons and serotonergic neurons in this circuit, we next crossed the URX/AQR/PQR ablation worms or BAG ablation worms into *C. elegans* with HIF-1 stabilized in the ADF neurons. Ablation of either the URX/PQR/AQR or the BAG neurons abrogated lifespan extension in the ADF HIF-1 stabilized worms. This suggests that oxygen sensing neurons act downstream of ADF HIF-1 stabilization in this pathway (**Fig. 5D-E**).

**Figure 5:**
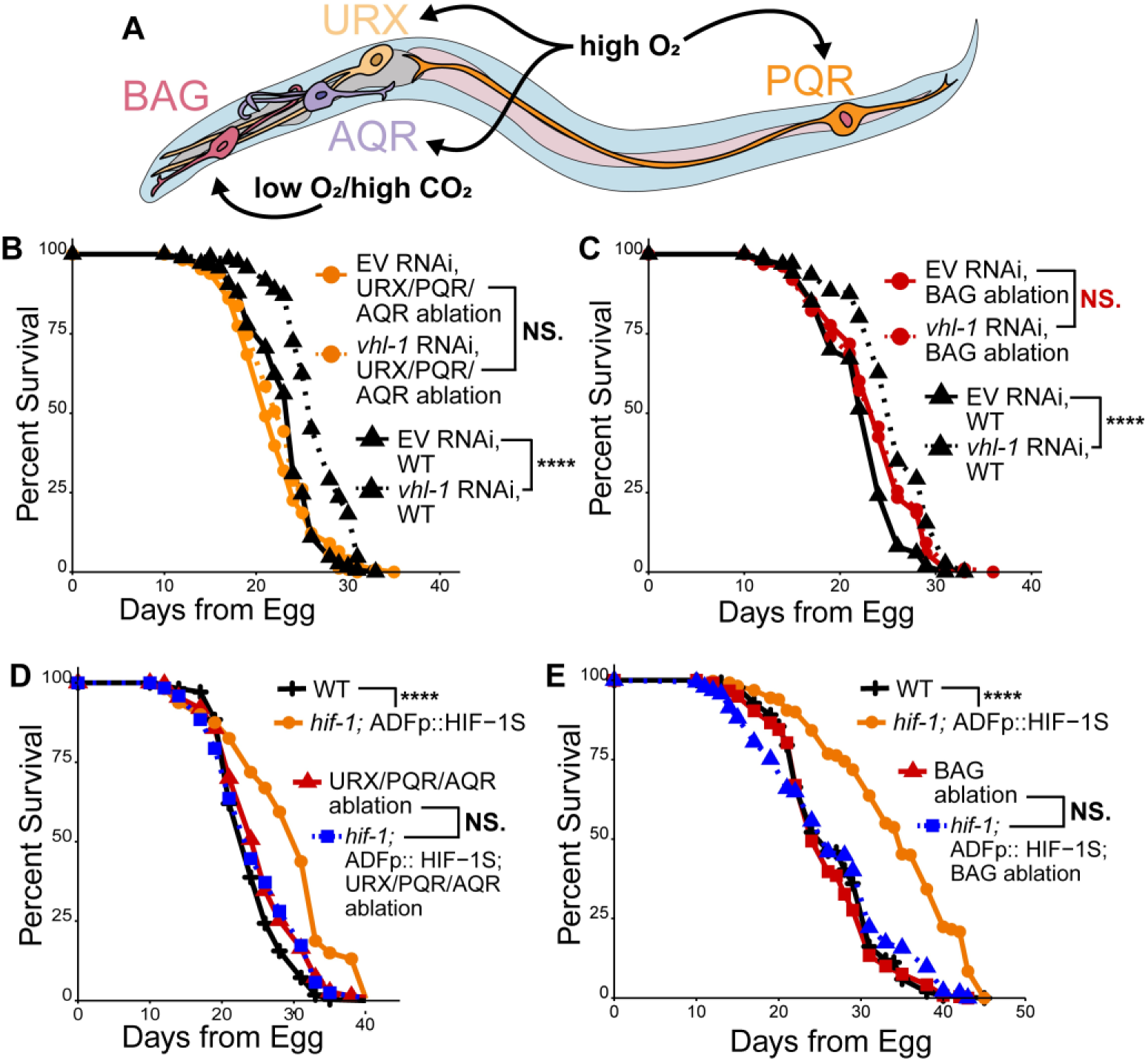
High- and low-oxygen sensing neurons are important for genetic activation of the hypoxic response to extend lifespan. (**A**) Diagram of oxygen sensing neurons in *C. elegans*. (**B**) Survival curves of WT and URX/PQR/AQR ablated worms (*qaIs2241 [gcy-36::egl-1 + gcy-35::GFP + lin-15(+)*]) on empty vector (EV) and *vhl-1* RNAi. *N* ≥ 279 worms per condition. (**C**) Survival curves of WT and BAG ablated worms (*Ex[gcy-31p::caspase-3(p12)-nz]; Ex[gcy-31p::cz-caspase-3 (p17)]; gcy-31p::GFP*) on empty vector (EV) and *vhl-1* RNAi. *N* ≥ 188 worms per condition. (**D**) Survival curves of WT, *hif-1(ia4);* ADF-HIF-1S, URX/PQR/AQR (*qaIs2241 [gcy-36::egl-1 + gcy-35::GFP + lin-15(+)*]) ablation, and URX/PQR/AQR (*qaIs2241 [gcy-36::egl-1 + gcy-35::GFP + lin-15(+)*]) ablation; *hif-1(ia4);* ADF-HIF-1S worms. *N* ≥ 150 worms per condition. (**E**) Survival curves of WT, *hif-1(ia4);* ADF-HIF-1S, BAG (*Ex[gcy-31p::caspase-3(p12)-nz]; Ex[gcy-31p::cz-caspase-3 (p17)]; gcy-31p::GFP*) ablation, and BAG (*Ex[gcy-31p::caspase-3(p12)-nz]; Ex[gcy-31p::cz-caspase-3 (p17)]; gcy-31p::GFP*) ablation; *hif-1(ia4);* ADF-HIF-1S worms. *N* ≥ 186 worms per condition. Significance in all panels is from a log-rank test comparing median survival. NS. = *p >* 0.05, * = *p* < 0.05, ** = *p* < 0.01, *** = *p* < 0.001, and **** = *p* < 0.0001. In all panels, three replicates were plotted together, and all statistics include a Bonferroni correction for multiple comparisons.

### The neuropeptide NLP-17 and its receptors contribute to *vhl-1-*mediated longevity

After identifying neurons, neurotransmitters, and neuroreceptors acting in the *vhl-1*-mediated longevity circuit, we wondered how this neural circuit ultimately propagates information about hypoxic conditions to the intestine and induces the pro-longevity gene *fmo-2*. The *C. elegans* nervous system does not directly innervate the intestine. Instead, neurosignaling molecules bind to receptors on peripheral tissues through packaging in dense-core vesicles and subsequent release into and diffusion through pseudocoelomic fluid^66,67^. These dense-core vesicles are packaged with a wide variety of neuropeptide signals, although there is also evidence that bioaminergic neurotransmitters like serotonin, dopamine, and adrenaline/noradrenaline can also signal through this dense-core vesicle mechanism^67–69^.

We first tested if inducing the hypoxic response through *vhl-1* RNAi extends lifespan in a mutant strain lacking the dense-core vesicle packaging gene *unc-31*. We observed that *unc-31* was required for *vhl-1* knockdown to extend lifespan (**Fig. 6A**). To examine whether neuropeptide signals carried by dense-core vesicles are involved, we knocked down two neuropeptide processing enzymes—*egl-3* and *egl-21*^70^. Successful knockdown was validated with qPCR (**Fig. S5A**). Interestingly, *egl-21* but not *egl-3* was required for *vhl-1-*mediated longevity (**Fig. 6B-C**). This could be because *egl-21* is the only enzyme known to perform the carboxypeptidase E cleavage step of neuropeptide processing^71^, while there are four known proprotein convertases with similar function to *egl-3*^70,72^.

**Figure 6.**
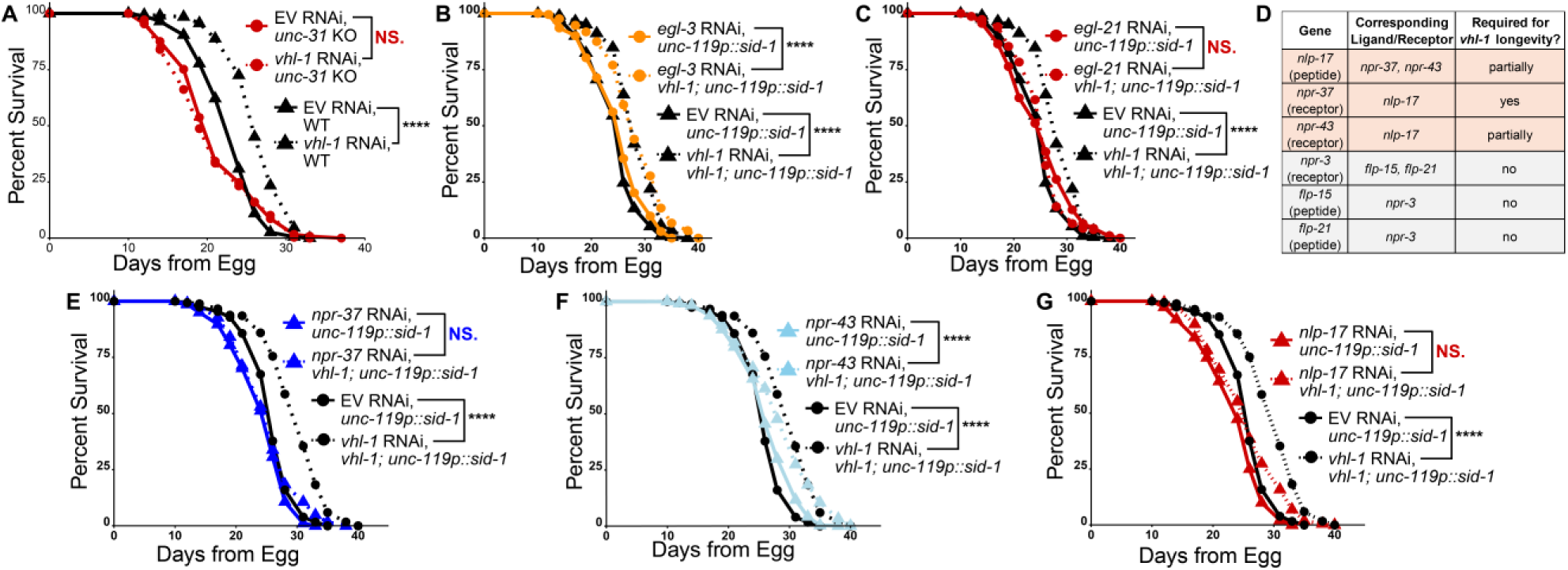
The neuropeptide *nlp-17* and its receptors *npr-37* and *npr-43* are required for *vhl-1-*mediated longevity. (**A**) Survival curves of *unc-31(e169)* and WT worms on empty vector (EV) and *vhl-1* RNAi. *N ≥* 242 worms per condition. (**B-C**) Survival curves of TU3311 (*unc-119p::sid-1)* and *vhl-1(ok161);* TU3311 (*unc-119p::sid-1)* worms on empty vector (EV) and *egl-3* (**B**) or *egl-21* (**C**) RNAi. *N ≥* 209 worms (**B**) and *N ≥* 185 worms (**C**) per condition. (**D**) Summary table of two sets of neuropeptide ligands/receptors that blunted *vhl-1-*mediated lifespan. (**E-G**) Survival curves of TU3311 (*unc-119p::sid-1)* and *vhl-1(ok161);* TU3311 (*unc-119p::sid-1)* worms on empty vector (EV) and *npr-37* (**E**), *npr-43* (**F**), or *nlp-17* (**G**) RNAi. *N ≥* 288 worms (**E**), *N ≥* 309 worms (**F**), and *N ≥* 309 worms (**G**) per condition. Significance in panels A-C and E-G are from log-rank test comparing median survival. Cox Regression for a significant interaction between *npr-43* knockdown and *vhl-1* knockout on lifespan, *p* <0.01, **. Cox regression for a significant interaction between *nlp-17* knockdown and *vhl-1* knockout on lifespan, p < 0.05, *. NS. = *p >* 0.05, * = *p* < 0.05, ** = *p* < 0.01, *** = *p* < 0.001, and **** = *p* < 0.0001. In all panels, three replicates were plotted together, and all statistics include a Bonferroni correction for multiple comparisons.

We next tested a group of 6 genes corresponding to neuropeptide/receptor binding pairs that blunted *vhl-1*-mediated *fmo-2* induction (**Fig. 6D**). These genes, along with their corresponding receptors, were targeted for RNAi and tested for lifespan in TU3311 (*unc-119p::sid-1*) and TU3311 (*unc-119p::sid-1*); *vhl-1 (ok161)* strains. We found that the *npr-3* receptor and its ligands, *flp-15* and *flp-21* were not required for *vhl-1*-mediated longevity. (**Fig. S5B-D**). However, knocking down the neuropeptide *nlp-17* partially blocked *vhl-1*-mediated longevity (**Fig. 6E**, Cox regression *p < 0.05*). Consistently, knocking down *nlp-17’*s receptors also completely (*npr-37*, **Fig. 6F**) or partially (*npr-43*, **Fig. 6G**, Cox regression *p < 0.001*) blocked *vhl-1-*mediated longevity. Successful knockdown of *npr-37* and *npr-43* was validated with qPCR (**Fig. S5A**). The *nlp-17* ligand result was validated using a genetic knockout (*nlp-17 (ok3461)*), which completely prevented *vhl-1* knockdown from extending lifespan (**Fig. S5E**). To test whether *nlp-17* acts downstream of other signals in this circuit, we measured the expression of *nlp-17* in ADF- and NSM-specific HIF-1 stabilized strains. Compared to WT and *hif-1* KO controls, we observed no change in *nlp-17* expression when HIF-1 was stabilized in the ADF or NSM serotonergic neurons (**Fig. S5F**). This could suggest either that *nlp-17* signaling acts in parallel to serotonergic signaling following genetic activation of the hypoxic response. Alternatively, neuronal HIF-1 stabilization may modify *nlp-17* splicing or translation without resulting in detectable differences in mRNA levels. Together, these data indicate NLP-17 signaling is required for longevity following genetic activation of the hypoxic response, although whether this peptide is synthesized or released in response to hypoxic response remains unclear.

**Supplemental Figure 5.**
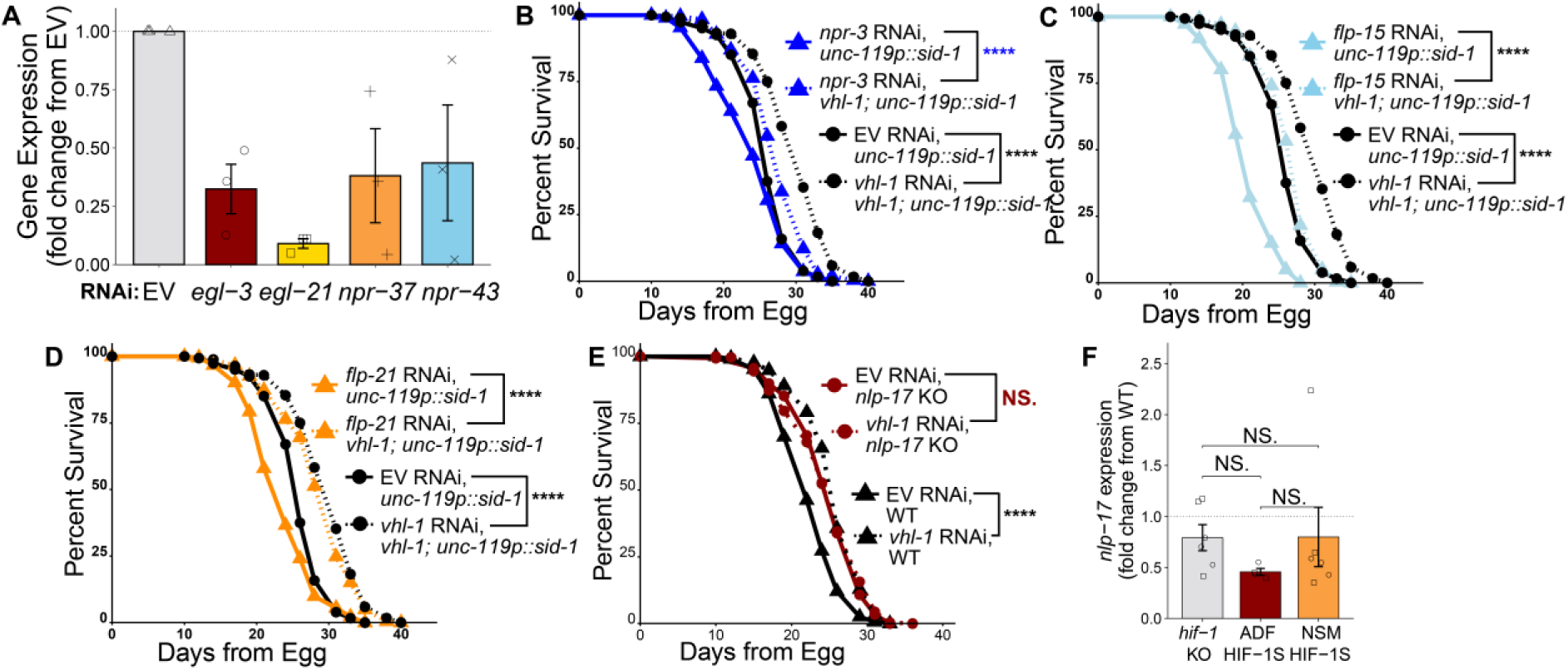
qPCR validation of RNAi hits from Figure 5, and neuropeptide screen hits that did not validate. (**A-B**) Gene expression of (**A**) *egl-3, egl-21, npr-37,* and *npr-43* in TU3311 (*unc-119p::sid-1)* worms raised on RNAi targeting each gene compared to control worms raised on empty vector (EV) RNAi for two generations. *N* ≥ 300 worms per replicate, or 900 worms per condition. The top of the bar represents the mean and error bars represent SEM. (**B-D**) Survival curves of TU3311 (*unc-119p::sid-1)* and *vhl-1(ok161);* TU3311 (*unc-119p::sid-1)* worms on empty vector (EV) and *npr-3* (**B**), *flp-15* (**C**), or *flp-21* (**D**) RNAi. *N ≥* 191 worms (**B**), *N ≥* 181 worms (**C**), and *N ≥* 183 worms (**D**) per condition. (**E**) Survival curves of *nlp-17(ok3461)* on empty vector (EV) or *vhl-1* RNAi. *N ≥* 275 worms per condition. (**F**) Gene expression of *nlp-17* in WT, *hif-1(ia4)* knockout, *hif-1(ia4); ADF:HIF-1S,* and *hif-1(ia4); NSM:HIF-1S* worms. *N* ≥ 200 worms per replicate, or 800 worms per condition. The top of the bar represents the mean and error bars represent SEM. Significance is from a two-tailed Wilcoxon rank sum test. Significance in panels B-E are from log-rank test comparing median survival. NS. = *p >* 0.05, * = *p* < 0.05, ** = *p* < 0.01, *** = *p* < 0.001, and **** = *p* < 0.0001. In all panels, 2-3 replicates were plotted together, and all statistics include a Bonferroni correction for multiple comparisons.

Taken together, these results support a model in which HIF-1 stabilization in the ADF serotonergic neurons leads to modified serotonin signaling from the ADF to the SER-7 expressing and GABA-producing RIS neuron (**Fig. 7A-B**). Oxygen sensing neurons act downstream of HIF-1 stabilization in the ADF neuron in this longevity circuit. Tyramine produced by the RIM neuron and neuropeptide (NLP-17) signaling are also required for genetic activation of the hypoxic response to extend lifespan, although it remains unclear whether these signals act upstream, downstream, or in parallel to the serotonergic and GABAergic components in this pathway.

**Figure 7.**
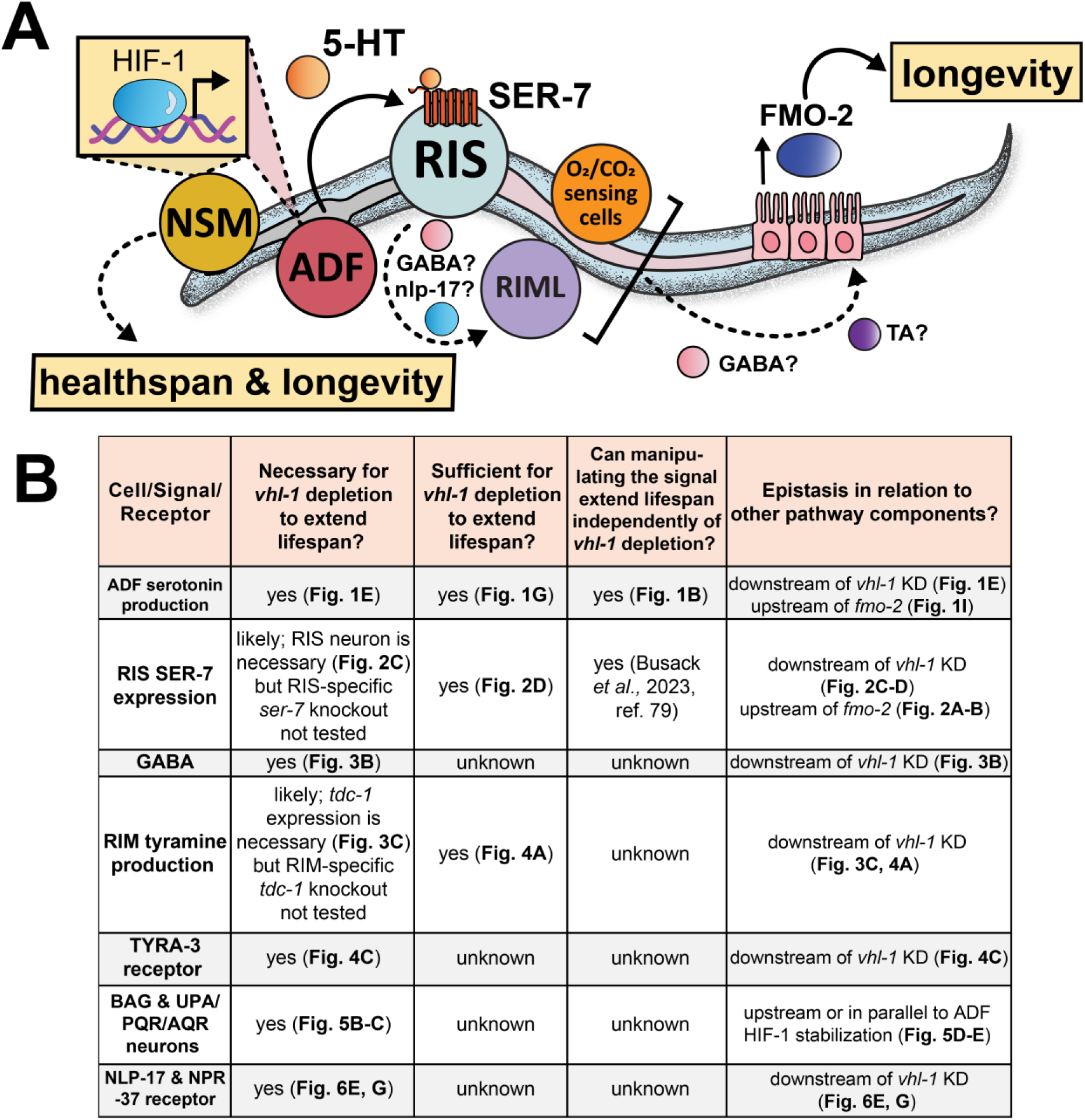
Working model of longevity following genetic activation of the hypoxic response. **A)** Schematic of working model. **B)** Table summarizing the necessity, sufficiency, and epistasis of neural signaling components of the working model.

## Discussion

In this study, we interrogated a complex signaling pathway through which genetic induction of the hypoxic response extends lifespan in *C. elegans*. Within this pathway, we demonstrate that: 1) stabilization of HIF-1 in the ADF serotonergic neurons modifies signaling to the SER-7 receptor on the GABAergic and neuropeptidergic RIS neuron (**Fig. 1-2**); 2) tyraminergic signaling through the RIM neuron and the TYRA-3 receptor (**Fig. 3-4**) are required for *vhl-1*-mediated longevity; 3) oxygen and carbon-dioxide sensory neurons act downstream of HIF-1 stabilization and serotonergic ADF neurons to extend lifespan (**Fig. 5**); and 4) in addition to neurotransmitter signals, the neuropeptide NLP-17 and its receptors NPR-37 and NPR-43 are required for genetic activation of the hypoxic response to extend lifespan (**Fig. 6**). Ultimately, these neural signals converge on the intestine, to activate FMO-2 and extend lifespan (working model summarized in **Fig. 7**). Evidence for the necessity, sufficiency, and epistatic relationships between each signal are summarized in **Fig. 7B**. In brief, all signaling molecules presented in this work are necessary for *vhl-1* to extend lifespan. Rescuing ADF serotonin production, RIS *ser-7* expression, and RIM tyramine production is sufficient for *vhl-1* to extend lifespan. The sufficiency of oxygen sensing neurons BAG and UPA/PQR/AQR, and the neuronal signals of GABA, NLP-17, and TYRA-3 to restore *vhl-1* mediated longevity remains unclear. All signals act downstream of *vhl-1*. ADF HIF-1 stabilization and SER-7 signaling act upstream of *fmo-2* induction, and the oxygen sensing neurons (BAG, UPA/PQR/AQR) act upstream of or in parallel to ADF HIF-1 stabilization.

The hypoxic response is highly conserved across species, highlighting the potential relevance of this pathway in other organisms, including humans. In nematodes, activating the hypoxic response promotes health and longevity. However, the physiological changes induced by the hypoxic response are broad and involve adaptations that can drive tumorigenesis in mammals. Therefore, a mechanistic understanding of the individual cells, signals, and circuits that mediate the beneficial effects of hypoxia is essential. In this work, the discovery that HIF-1 stabilization in the ADF neurons extends lifespan by 26% demonstrates the potential for targeted manipulations within this pathway to have large effects on lifespan. Additionally, our data identify a previously unknown role for tyramine/adrenaline in genetic activation of the hypoxic response as well as establish NLP-17, a neuropeptide with no previous known function, as a signal in *vhl-1-*mediated longevity. This work also demonstrates that the oxygen sensing AQR, PQR, URX, and BAG neurons act downstream of the serotonergic ADF neuron in genetic activation of the hypoxic response, indicating an interaction between internal and external oxygen sensing mechanisms in longevity. Together, the mapping of these cells, signaling molecules, and receptors acting downstream of genetic activation of the hypoxic response lays the foundation to discover druggable targets that selectively modulate aging in humans without negative side effects.

Other studies in *C. elegans* have also identified neural networks involving the hypoxic response. For example, previous work found a role for HIF-1 in hypoxia-mediated behaviors, such as food-dependent hyperoxia (excessive oxygen) avoidance^73^. Hyperoxia avoidance requires *hif-1* expression both in the neurons and the tyraminergic uv1 neuroendocrine cells. Interestingly, serotonin signaling from the ADF neuron is also required for WT-like hyperoxia avoidance^73^. While this HIF-1-mediated behavioral circuit has parallels with the HIF-1-mediated longevity circuit interrogated in this work, these pathways do diverge: the key tyraminergic cell type implicated in hyperoxia avoidance is the uv1 gonadal neuroendocrine cells, while the RIM tyraminergic neurons played a role in longevity (**Fig. 4**). Together, the partial overlap between this behavioral and longevity circuit implies that modifying neural signaling to induce the hypoxic response and extend lifespan may also alter behavioral phenotypes. Future research in this area should interrogate exactly where these and any other HIF-1-mediated circuits overlap and diverge.

While this study identifies many neural signals required for *vhl-1* knockdown or knockout to extend lifespan, one key limitation of this work is the potential differences between genetic and environmental methods of inducing the hypoxic response. While *vhl-1* knockdown or knockout leads to HIF-1 stabilization by blocking its proteasomal degradation, it also results in hydroxylated but stable HIF-1. This contrasts with environmental hypoxia, in which HIF-1 remains stable because it cannot be hydroxylated. While HIF-1 is stabilized and localized to the nucleus in both cases, there are differences in transcriptional outcomes between stable hydroxylated and unhydroxylated states^74,75^. Additionally, we did not explore the effects of alternative genetic activators of the hypoxic response such as HIF-1 hydroxylase PHD/EGL mutants, which may provide further insight into how different manipulations of the hypoxic response impact longevity. Future work should interrogate the similarities and differences between the circuits driving longevity in response to environmental hypoxia, *vhl-1* knockdown, HIF-1 stabilization, and PHD/EGL knockdown.

The circuit-mapping approaches employed in this work are also impacted by limitations in cell-specific genetic modifications and in the use of RNAi knockdown. For example, cell-specific rescue constructs can sometimes lead to unintended rescues in other cell types due to cell-nonautonomous signaling. Because all serotonin-producing neurons also express the serotonin reuptake transporter *mod-5*, serotonin produced by one cell in our rescue strains could be taken up by other serotonin-producing neurons, leading to unintended signaling effects. This may also be true of the uv1 and RIM tyraminergic rescue strains, although little is known about tyramine reuptake in *C. elegans*. Two cells identified via tissue-specific rescue experiments (ADF and RIS, **Fig. 1G** and **Fig. 2D**) were also found to be necessary via cell-specific knockout (ADF *tph-1* KO, **Fig. 1E**; RIS ablation, **Fig. 2C**), decreasing the likelihood of a false positive from the rescue strain technique. However, the role of the RIM neuron was identified via a *tdc-1* rescue strain and was not validated using a RIM-specific knockout (**Fig. 7B**). Therefore, the contribution of RIM signaling to this circuit is less well-validated, and a RIM-specific *tdc-1* knockout strain should be examined in future work. Finally, additional genetic controls could better support the role of RIS*-*specific *ser-7* expression in genetic activation of the hypoxic response. For example, a *ser-7* expressing neuron that was not a hit in our screen could also be ablated and tested for necessity in *vhl-1* mediated longevity. This experiment would test whether the ability of RIS ablation to attenuate *vhl-1*-mediated longevity is not a false positive driven by any disruption to *ser-7* expression.

Additionally, although RNAi experiments were validated using sequence confirmation and qPCR, not all RNAi hits were corroborated with genetic knockouts. This leaves open the possibility that RNAi-induced knockdown effects differ from complete genetic ablation, or that production of a given RNAi may modify bacterial metabolism in a way that indirectly modifies the hypoxic response in *C. elegans*. Finally, while the use of

RNAi knockdown and genetic knockouts establishes the necessity of many signals within the *vhl-1*-mediated longevity circuit, the exact directionality of these signals remains unclear. It is possible that increased, decreased, or pulsatile changes in signaling through these bioamines and neuropeptides are required for genetic activation of the hypoxic response to extend lifespan. Work on *C. elegans* reversal behavior has also revealed an antagonistic relationship between RIM and RIS activity facilitated by both chemical (neuropeptide and tyramine) and electrical (gap junction) signaling^76,77^. This known interaction should also be interrogated in the context of how these cells may communicate following genetic induction of the hypoxic response. Future work in this area could use tools to measure or modify neuronal activity, such as calcium imaging or optogenetics, to begin answering these questions.

While many individual neurosignaling components are essential for genetic activation of the hypoxic response to extend lifespan, their epistasis is unclear (**Fig. 7B**). Most components of the pathway identified in this work act downstream of *vhl-1* depletion, and upstream of *fmo-2* induction (summarized in **Fig. 7A-B**). However, the order of each signal between these two endpoints is only predicted based on *C. elegans* neural wiring and the overlap between various identified signals and cells. For example, we hypothesize in our working model that GABA may be produced by the RIS neuron in this circuit because RIS is the primary GABAergic neuron required for *vhl-1*-mediated longevity. Alternatively, it is possible that GABA is produced by a different cell that either acts in series or in parallel with RIS signaling. In order to determine the order of each signaling component, future studies should generate genetic manipulations to each signaling component that may mimic *vhl-1* knockout to promote longevity, cross these new strains into knockouts of other signals required for *vhl-1* mediated longevity; and measure the lifespans of each double and triple mutant. This approach would also narrow down which signals are downstream of the genetic activation of the hypoxic response, and which are sufficient to extend lifespan upstream of the hypoxic response in a normoxic environment. One notable target for further exploration is the SER-7 expressing RIS neuron, which plays a role in sleep^77^ and stress resistance^78^, and can extend lifespan when optogenetically activated under normoxic conditions^79^.

Another limitation of this work is that it uses lifespan as the main readout for organismal health. While genetic manipulations that extend lifespan often improve stress resistance and healthspan^8,80^, longevity manipulations can also have adverse effects on reproduction^81,82^ and behavior^83,84^. In this study, we find that HIF-1 stabilization in the ADF neurons does not prevent the deleterious effects of *hif-1* knockout on mobility, but that HIF-1 stabilization in the NSM neurons may attenuate age-related decline in pumping and thrashing (**Fig. S1)**. However, future work should examine whether other modifications to this pathway, such as manipulations to RIM, RIS, or NLP-17 signaling, influence healthspan in addition to lifespan. It will be important for future studies to determine whether various components of this pathway affect both longevity and the response to different types of stressors like oxidative stress, proteotoxic stress, and infection, as HIF-1 activity also interacts with multiple stress responses^41,42 39,40,43^.

While our understanding of this circuit is still incomplete, our findings point to several promising signaling components that could be targeted for longevity interventions. Manipulating serotonin and tyramine/adrenaline signaling pathways may offer strategies for extending lifespan, with minimal pleiotropic effects if precisely controlled. These neural targets have great potential for longevity therapeutics due to 1) the small number of neurons that can control aging-related pathways in the entire organism; 2) the availability of FDA-approved pharmaceuticals that target individual bioaminergic transporters and receptors^85–87^; and 3) the high conservation of neurotransmitter biology between invertebrates and mammals^58,70,88^. Further research into how these pathways can be modulated in a targeted manner could spur development of interventions that promote healthy aging and delay the onset of age-related diseases.

## Methods

### Strains and Growth Conditions

*C. elegans* were cultured according to established protocols^31^. In brief, worms were grown at 20 °C on standard solid nematode growth media (NGM). Throughout their lifespan, the worms were fed *E. coli* OP50, except during RNA interference (RNAi) experiments, where *E. coli* HT115 was used to deliver double-stranded RNA. Transfers of worms were carried out using a platinum wire, unless stated otherwise. The RNAi strains employed are listed in Supplementary Table 1, while the strains used in the experiments are provided in Supplementary Table 2. Genotypes were verified through PCR, and RNAi imaging results were confirmed by sequencing and quantitative PCR (qPCR) before proceeding with the experiments.

### Generating transgenic strains

#### Neuron-specific stabilized HIF-1 strains

We used the following promoters to drive cDNA of HIF-1 (P621A)::SL2::GFP in three serotonergic neuronal populations: ADF neurons (*srh-142p::* HIF-1 (P621A)), NSM neurons (*ceh-2p::* HIF-1 (P621A)), HSN neurons (*ham-2p::* HIF-1 (P621A)). All plasmids were verified via restriction digest and sanger sequencing, and ApE files are available upon request. Plasmids were microinjected to *hif-1(ia4)* KO strain by Suny Biotech using the co-injection marker *myo-2p*::GFP and 2-3 transgenic lines were tested in each experiment.

#### Neuron-specific *tph-1* rescues

The pKA805[*srh-142p*::TPH-1] and pKA807[*ceh-2p::*TPH-1] constructs were generously provided by Dr. Kaveh Ashrafi. These constructs were injected at ∼50 ng/µL) with fluorescent co-injection marker *myo-2p*::mNeonGreen (15 ng/µL) or *sur-5p*::*sur-5::*NLSGFP (20 ng/µL) and junk DNA (up to 100 ng/µL) into gonads of day 1 gravid adult hermaphrodites. Standard protocols were followed to isolate and obtain stable over-expression mutants^89^.

#### *ser-7* rescue strains

Plasmid construction and microinjection was conducted by Suny Biotech to generate all *ser-7* rescue constructs. In designing the plasmids, we used the following promoters to drive cDNA of *ser-7*::SL2::GFP in different neuronal populations: full rescue (*ser-7p::ser-7)*, interneuron rescue (*glr-1p::ser-7)*, bioaminergic neuron rescue (*cat-1p::ser-7)*, glutamatergic neuron rescue (*eat-4p::ser-7*), GABAergic neuron rescue (*unc-47p::ser-7*), GABAergic motor neuron rescue(*unc-25p::ser-7*), sensory neuron rescue (*osm-6p::ser-7*), cholinergic motor neuron rescue(*acr-2p::ser-7*), cholinergic neuron rescue (*unc-17p::ser-7*), M3 & M4 neuron rescue (*ceh-28p::ser-7*), and intestinal rescue (*vha-6p::ser-7*). All plasmids were verified via restriction digest and sanger sequencing, and ApE files are available upon request. Plasmids were microinjected by Suny Biotech using the co-injection marker *myo-2p::*GFP and 2-3 transgenic lines were tested in each experiment.

#### RIS, RIC, and BAG neuronal ablation strains

For RIS neuronal ablation strains, we purchased donor plasmid *mec-18p*::caspase-3 (p12)::nz [TU#813] (Plasmid #16082) and *mec-18p* cz::caspase-3 (p17) [TU#814] from Addgene (Plasmid #16083), and used Gibson cloning to replace *mec-18p* with *srsx-18p*. Three constructs *srsx-18p*::caspase-3(p12)::nz, *srsx-18p*::cz::caspase-3(p17), and *srsx-18p*::GFP were co-injected with fluorescent co-injection marker *myo-3p*::GFP (20 ng/µL) into the wild-type N2 strain to generate RIS genetic ablation strains. Similarly, for RIC ablation strain, *tbh-1p* was constructed into TU#813 and TU#814 to replace *mec-18p*. Three constructs *tbh-1p*::caspase-3(p12)::nz, *tbh-1p*::cz::caspase-3(p17), and *tbh-1p*::GFP were co-injected with fluorescent co-injection marker *myo-3p*::GFP (20 ng/µL) into the wild-type N2 strain to generate RIC genetic ablation strains. For BAG ablation strain, *gcy-31p* was constructed into TU#813 and TU#814 to replace *mec-18p*. Three constructs *gcy-31p*::caspase-3(p12)::nz, *gcy-31p*::cz::caspase-3(p17), and *gcy-31p*::GFP were co-injected with fluorescent co-injection marker *myo-3p*::GFP (20 ng/µL) into the wild-type N2 strain to generate BAG genetic ablation strains. All plasmids were verified via restriction digest and sanger sequencing. ApE files available upon request. Plasmid construction and microinjection was conducted by Suny Biotech.

#### RIM and uv1 *tdc-1* rescue strains

Plasmid construction and microinjection was conducted by Suny Biotech to generate both *tdc-1* rescue constructs. In designing the plasmids, we used the *ocr-4* promoter to drive cDNA of *tdc-1*::SL2::GFP in the uv1 neuroendocrine cells. To express cDNA of *tdc-1*::SL2::GFP in the RIML neuron, we used the *gcy-13* promoter. All plasmids were verified via restriction digest and sanger sequencing, and ApE files are available upon request. Plasmids were microinjected by Suny Biotech using the co-injection marker *myo-2*p::GFP.

### RNAi Knockdown

For all RNAi knockdowns, worms were exposed to the RNAi treatment for two generations to achieve optimal knockdown. All RNAi clones were sourced from the Vidal RNAi library. Each RNAi clone was sequence-verified.

### Quantitative PCR

For RNAi validation experiments, RNA was isolated from day 1 adult worms that had been grown on RNAi for two generations. For measuring *fmo-2* induction in the *hif-1(ia4);* ADF::HIF-1S strain, worms were grown on OP50, synchronized, and collected at day 1 of adulthood. RNA extraction was performed using the Direct-zol RNA Miniprep Kit (Zymo), and cDNA synthesis was carried out with the Biorad iScript cDNA Synthesis Kit. Gene expression was assessed using the SYBR Green quantitative RT-PCR ^90^ system (BioRad), with mRNA levels normalized to the housekeeping genes *cdc-42* and *Y45FD10.4*. Primers for RNAi validation were designed to target the 3’ UTR of each gene to avoid amplification of the bacterially produced RNAi. Gene expression was quantified using standard curves.

### Lifespan Measurements

Lifespan assays were conducted following previously described protocols^31^. Briefly, 10-15 gravid adults were transferred to NGM plates for a three-hour timed egg lay, after which they were removed. Once the progeny reached day 1 of adulthood, 60-80 worms were placed on NGM plates containing 33 µL of 150 mM fluorodeoxyuridine (FUdR) and 100 µL of 50 mg/mL ampicillin per 100 mL of NGM. FUdR inhibits progeny development, while ampicillin prevents bacterial contamination. NGM + FUdR + Ampicillin plates were seeded with concentrated bacteria (5x for OP50-fed lifespan assays). At least two plates per strain and condition were used for each lifespan replicate. Worms were considered dead when they failed to respond to a gentle touch with a platinum wire under a dissection microscope. Lifespan data were recorded at least three times a week until all worms were dead. To prevent escape, a barrier of 75 µL of 100 mM palmitic acid (Sigma-Aldrich) dissolved in 100% ethanol was applied along the edges of each lifespan plate. Data were analyzed using R version 4.3.1 and visualized in Adobe Illustrator 2022.

### RNAi Lifespans

RNAi lifespan assays were performed similarly to standard lifespan assays, with modifications to the initial timed egg lay (TEL) procedure and food concentrations. To ensure maximal RNAi knockdown, worms were first TEL’ed for 3 hours on RNAi plates. The progeny from this TEL were left on RNAi plates to develop into gravid adults and then used for a second TEL under the same RNAi conditions. Progeny from the second-generation TEL were then used for the lifespan assay. RNAi bacteria (HT115) are ampicillin-resistant and can grow slowly on AMP-containing lifespan plates. To maintain consistent bacterial availability throughout the lifespan assay, RNAi plates were seeded with bacteria at 2x concentration, starting from an optical density (OD_600_) of 3.0.

### Fluorescent Slide Microscopy

Fluorescent images for this study were captured using a Leica M165F fluorescent microscope, controlled by Leica Application Suite X (LASX) software. A minimum of 15 worms per condition were imaged at a magnification of at least 70x. For image quantification, individual worms were carefully separated to ensure they did not touch. Custom R code^91^ was then used to create a pixel mask of each worm from the brightfield image and measure the fluorescent intensity of that region in the corresponding fluorescent image. The background fluorescence was subtracted from the mean fluorescence values. Data were analyzed using R version 4.3.1 and visualized in Adobe Illustrator 2022.

### Pumping Assays

Pumping was measured in day 2 gravid adults, as previously described^31^. The number of pumps per 30 seconds was measured in at least 10 worms under 100x magnification. Because food availability is known to change pumping rate, only worms on the bacterial lawn were considered in this assay. Data were plotted by R version 4.3.1 and Adobe Illustrator 2022.

### Thrashing Assays

Thrashing was measured in day 2 gravid adults, as previously described^31^. Briefly, approximately 10 worms were placed in a droplet of M9 solution. Once worms began moving at a maximum rate, their body bends were counted for 30 seconds. Data were plotted by R version 4.3.1 and Adobe Illustrator 2022.

### 30-minute Video Recordings for Velocity and Movement Patterns

On day 1 and day 5 of adulthood, 10-15 worms of each genotype were placed in the center of a 35 mm petri dish that was completely covered in 350 μL of OP50 bacteria (bacteria was grown overnight to an optical density (OD_600_ 3.0). Worms were placed above the bright light source at the base of the microscope used for video recording for 10 minutes to acclimate to the light, and then 30-minute video recordings were started. Videos were taken at 3.65x magnification, and at 1 frame per second on a Leica M165F microscope using Leica Application Suite X (LASX) software and 4×4 binning. The videos were analyzed using the wrMTrck multiple object tracker plugin (wrMTrck_Batch v1.04) on FIJI^92^ (FIJI Is Just ImageJ bundled with 64-bit Java 1.8.0) to measure average speed, maximum speed, worm area, total distance traveled, and distance from the point of origin. Not all worm paths lasted the full 30-minute duration of the video due to fleeing and collisions. In cases when worms fled the plate and were no longer visible during the video, the data from that worm was included if the on-plate duration was ≥10 minutes. In cases where worms collided and wrMTrck failed to correctly determine which worm was which after the event, the data for those worms before the collision was included if the duration was ≥ 10 minutes. In all data analysis using wrMTrck data, we confirmed that there was no significant difference in average path duration per condition. Data were plotted by R version 4.3.1 and Adobe Illustrator 2022.

### Statistical Analysis

The height of all bar plots represents the mean value for each condition, with error bars indicating the standard error of the mean (SEM). For comparisons involving more than two conditions, one-way ANOVA followed by post-hoc Tukey HSD tests ^47^ were used to assess interactions between variables and determine statistical differences between conditions. Significance levels are indicated as *p < 0.05, **p < 0.01, ***p < 0.001, and ****p < 0.0001.

For lifespan assays, statistical comparisons of survivorship curves were performed using the survfit survival analysis function in R. The log-rank test was used to compare two survival curves. To assess interactions across multiple experimental variables and for comparisons involving more than two survival curves, a Cox regression analysis was conducted using the survivalMPL package in R. For experiments with multiple biological replicates, p-values were adjusted using the Bonferroni correction. Exact sample sizes (Ns) for each experiment are provided in the source data files, and minimum sample sizes for each plot are specified in the figure legends.

## Supporting information

Supplementary Table 1, RNAi

Supplementary Table 2, Strains

